# Spiny mouse fibroblasts exhibit a baseline preference for glycolysis and are resilient to oxidative stress across lifespan

**DOI:** 10.1101/2025.10.02.680123

**Authors:** Ebenezer Aryee, Ajoy Aloysius, Sandeep Saxena, Renee Donahue, Brennan Riddell, Hemendra Vekaria, Patrick G. Sullivan, Huazhen Liu, Pradeep Kachroo, Ashley W. Seifert

**Affiliations:** Department of Biology, University of Kentucky, Lexington, KY 40508, USA; Aligning Science Across Parkinson’s (ASAP) Collaborative Research Network, Chevy Chase, MD, 20815, USA; Department of Neuroscience, University of Kentucky, Lexington, KY 40506, USA; The Spinal Cord and Brain Injury Research Center (SCoBIRC), University of Kentucky, Lexington, KY 40506, USA; Lexington VA Medical Center, Lexington, KY 40506, USA; Department of Plant Pathology, University of Kentucky, Lexington, KY 40546, USA

**Keywords:** mitochondria, mitochondrial metabolism, electron transport system, reactive oxygen species, regeneration, aging

## Abstract

Although metabolism was originally studied across an array of mammals, contemporary metabolic studies moved towards human and mouse cells. A newfound interest in how metabolism regulates cell state during homeostasis, tissue repair and disease has uncovered key roles for energy flux inside and outside of mitochondria. Fibroblasts are key mediators of wound healing outcomes and prior work uncovered that cells from highly regenerative mammals (spiny mice and rabbits) exhibit enhanced resistance to oxidative stress compared to those from non-regenerating laboratory mice and rats. Using a battery of cellular tests in primary ear pinna fibroblasts from spiny mice, rabbits, laboratory mice and rats, we show that cells from spiny mice and rabbits exhibit a baseline preference for glycolysis supporting lower ROS-production. Mitochondria from spiny mouse fibroblasts were generally low respiring and depolarized while exhibiting a large, spherical morphology. We observed this large, spherical phenotype consistently across lifespan in ear pinna fibroblasts from fetal, young and old spiny mice and cells from all ages were highly resistant to oxidative stress. While rabbit, mouse and rat fibroblasts had polarized tubular mitochondrial networks typical of adult mammalian fibroblasts, isolated rabbit and spiny mice fibroblasts shared lower oxygen consumption efficiency even in the absence of a potential gradient. Taken together, our results support that a shared metabolic signature exists in stromal cells from highly regenerative mammals, although possibly driven by different mechanisms, to converge on a ROS-resistant phenotype that increases cellular resilience.

## INTRODUCTION

It is widely accepted that most mammals repair damaged skin and musculoskeletal tissues with a scar (aka fibrotic repair). In response to tissue damage, a stereotypical sequence of events is initiated beginning with hemostasis and inflammation (1). Shortly thereafter, keratinocyte migration over the injury site restores barrier function (re-epithelialization) and a provisional extracellular matrix (ECM) is produced by cells in the wound site. Ultimately, this collagen-rich matrix is remodeled into mature scar tissue, and this overlapping sequence of events is the normal outcome of tissue damage in laboratory rodents and humans. In contrast, some vertebrates can functionally reconstruct injured tissues via reparative regeneration. While vertebrate regeneration typically combines processes observed during fibrotic repair and embryonic development (2, 3), key mechanisms that predispose how cells react to injury via regeneration or fibrotic repair remain obscure.

Spiny mice (*Acomys spp*.), have emerged as an important homeothermic animal model to investigate complex tissue regeneration (2). The regenerative abilities of *Acomys kempi* and *Acomys percivali* were first discovered as a response to full-thickness injuries to the skin and ear pinna with restoration of dermis, hair follicles, sweat glands and cartilage instead of scar tissue (4). Studies using *Acomys dimidiatus*, *A. kempi* and *Oryctolagus cuniculus* (New Zealand white rabbit) showed that ear pinna regeneration in these mammals was indeed a robust phenotype (5, 6) supporting previous experiments with rabbits (7–10). Subsequent research in regenerating (spiny mice, brush-furred mice) and non-regenerating rodents revealed that enhanced regenerative ability is confined to deomyine rodents (11). This phylogenetic restriction suggests specific cellular features may have evolved in these species which support enhanced regenerative ability.

Complex tissue regeneration requires accumulation of resident cells that proliferate and expand prior to cell differentiation; a transitional period referred to as blastema formation (12, 13). Importantly, what distinguishes cells transiting through a blastema from those that form a tissue analgen during development is that cells in the wounded tissue experience high oxidative stress from reactive oxygen species (ROS) and exposure to inflammatory cells and their products. Collectively, cells strive to maintain tissue and organismal homeostasis in the face of injury, disease and infection. Thus, cells act as frontline sensors to detect and mitigate extrinsic stressors (inflammatory, oxidative, etc.). The sensing capacity of cells depends, in part, on a basal metabolic state influenced by mitochondrial function which is highly sensitive to deviations in biomolecules between various cell compartments (14). Injury disrupts the local tissue microenvironment by altering oxygen and metabolite concentrations, which in turn affects conditions such as pH and ion flux. These alterations upset cellular equilibrium and activate the cell damage response (CDR), triggering changes in cellular behavior (15, 16). Therefore, cell autonomous features governed by mitochondrial and metabolic states can regulate genomic changes and direct resident cells to carry out a regenerative response (17, 18).

A connection between metabolic state and cell proliferation was better appreciated from observing the Warburg effect in cancer cells, where in the presence of ample amounts of oxygen, glucose breakdown through glycolysis shifted towards lactate production, avoiding carbon loss through TCA-produced CO_2_ (19). Specifically, halfway through glycolysis, intermediates can be shunted towards the synthesis of biomolecules like nucleotides and other biomass building blocks, as well as NADPH via the pentose phosphate pathway to support cell proliferation as seen in cancer cells and activated T cells (20). In addition to other reactions, NADPH synthesized from this pathway is utilized in redox balancing (21) and biomolecule synthesis to support cell proliferation (cholesterol, tetrahydrofolate, proline, fatty acids and deoxyribonucleotides) (22). A similar observation has been noted for cells participating in tissue regeneration (23, 24) suggesting metabolic reprogramming of connective tissue fibroblasts may be a key feature of vertebrate regeneration. Beyond cell proliferation, these distinctive metabolic signatures are equally pronounced in embryonic stem cells (ESCs). Compared to differentiated cells, ESCs are more dependent on glycolysis compared to oxidative phosphorylation (OxPhos) (25, 26). The correlation between cell potency, proliferation and glycolytic bias becomes more intimately linked in the case of cellular reprogramming, where reprogramming efficiency of fibroblasts to induced pluripotent stem cells (iPSCs) is linked to a pro-glycolytic metabolic state, with efficiency increased and decreased significantly in response to glycolytic inhibitors and augmenters respectively (27, 28).

Fibroblasts mediate the last three stages of wound repair (rev. in 29) and comprise the major proliferative cell type during regeneration (30, 31). This raises an important question: do cell autonomous mechanisms governed by cellular metabolic state regulate how fibroblasts respond to injury? Compared to mouse (*Mus musculus*) fibroblasts, *Acomys* fibroblasts show increased resistance to TGFβ1-induced myofibroblast activation (32). Additionally, *Acomys* and rabbit (*Oryctolagus cuniculus* – another highly regenerative mammal) fibroblasts exhibit enhanced resistance to radiation (up to 30 Gy) and ROS-induced senescence (up to 300µM) compared to fibroblasts obtained from non-regenerators; factors attributed in part to a rapid antioxidant response (18). These *in vitro* studies support the idea that intrinsic cellular mechanisms utilizing different signaling pathways, redox stabilizing reactions and possibly metabolic and epigenetic controls put *Acomys* fibroblasts in a state that primes them towards a regenerative response even in the absence of external factors such as tissue mechanics and inflammation. Since ROS generation and management is intimately linked to cellular metabolism, we hypothesized that stress resistant *Acomys* and *Oryctolagus* fibroblasts would possess a pro-glycolytic profile linked to mitochondrial physiology.

Here, we discover that primary ear pinna fibroblasts from *Acomys* and *Oryctolagus* show a pro-glycolytic baseline metabolic state compared to *Mus* and *Rattus* fibroblasts, which instead exhibit comparatively higher OxPhos states typical of differentiated fibroblasts. The pro-glycolytic mitochondrial state is also marked by higher glucose sensitivity. We also find that *Acomys* mitochondria present a unique phenotype being large, spherical and mostly depolarized. Lastly, examining *Acomys* primary ear pinna fibroblasts we find that this unique mitochondrial phenotype persists across lifespan and that cells retain their high resistance to oxidative stress independent of life-stage. Together, these results suggest that evolutionarily distant mammals may share a metabolic cell state reached via different mechanisms that can support regenerative healing and support the need to expand the range of mammalian models in biomedical research beyond the well-utilized few; especially in the field of wound healing.

## RESULTS

### Fibroblasts from regenerating mammals exhibit a relatively low basal metabolic rate and high glycolytic rate

Based on previous data demonstrating strong resistance to oxidative stress in *Acomys* and *Oryctolagus* ear pinna fibroblasts (18), we hypothesized this result could reflect a reduced metabolic state not typically observed in mouse and human fibroblasts. We ran conventional Seahorse Mitochondria Stress and Glycolytic Stress tests (MST and GST) using primary ear pinna fibroblasts from highly regenerative (*Acomys*, *Oryctolagus*) and non-regenerative (*Rattus*, *Mus*) mammals cultured for two passages (see Methods).

Using sequential injections of oligomycin, FCCP, and rotenone/antimycin A to inhibit ATP synthase, uncouple the mitochondrial proton gradient and maximize oxygen consumption, and inhibit complex I/III activity respectively, we measured oxygen consumption rates and calculated basal respiration, maximal respiration, and specific respiratory capacity (Fig. 1a-c and Supp. Fig. 1a-c). *Mus* fibroblasts exhibited significantly higher oxygen consumption at baseline compared to *Acomys*, *Rattus* and *Oryctolagus* fibroblasts (Fig. 1a-b). Despite having a low basal metabolic rate, in response to FCCP *Oryctolagus* fibroblasts rapidly increased oxygen consumption exhibiting maximal respiration levels matching that observed in *Mus* (Fig. 1a-c). *Oryctolagus* fibroblasts also exhibited high spare respiratory capacity levels that matched both *Rattus* and *Mus* (Supp. Fig. S1a). In contrast, maximal respiration and specific respiratory capacity (maximal respiration – basal respiration) were significantly lower in *Acomys* compared to the other species (Fig. 1a-c and Supp. Fig. S1a). ATP production and proton leak were significantly higher in *Mus* compared to the three other species (Supp. Fig. S1b-c).

**Figure 1.**
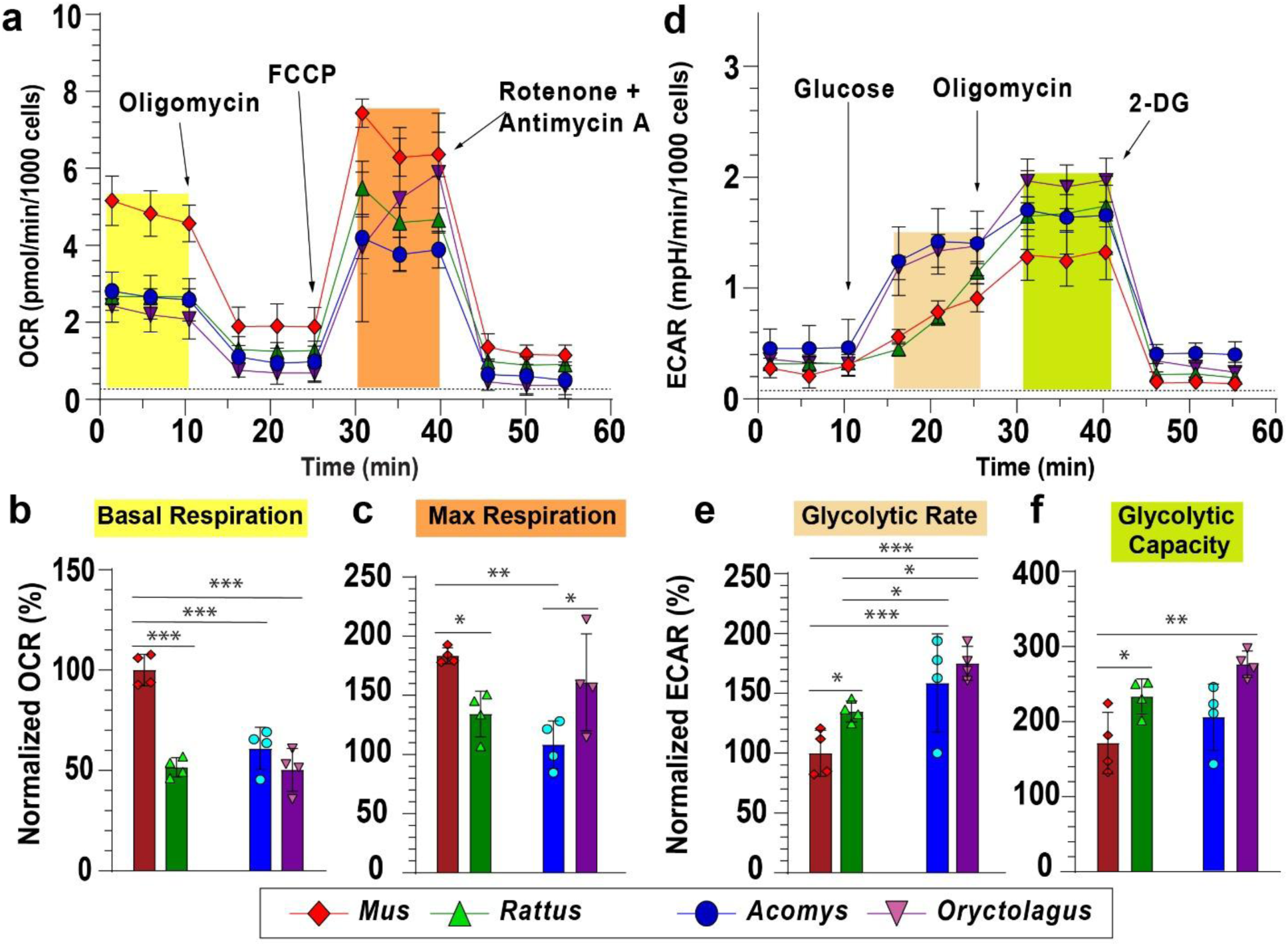
Fibroblasts from regenerating mammals exhibit a high glycolytic rate. MS-mitochondrial stress tests **(a-c)** and GST-Glycolytic tests **(d-f)** run on Seahorse XFe96 analyzer using primary ear pinna fibroblasts from *Acomys*, *Oryctolagus*, *Mus* and *Rattus*. Oxygen consumption rates (OCR) and extracellular acidification rates (ECAR) were normalized to the basal respiration and glycolytic rates in *Mus*. Basal respiration levels were significantly higher in *Mus* compared to the other three species (ANOVA, *F*=28.1749, *p*<0.0001) (a-b). Maximal respiration was significantly different among species (ANOVA, *F*=6.775, *p*=0.0063), with higher rates in *Mus* compared to *Acomys* and *Rattus* but not *Oryctolagus* (*p* = 0.2286) (c). Maximal respiration in *Oryctolagus* was significantly higher compared to *Acomys*. Fibroblasts from *Acomys* and *Oryctolagus* exhibited significantly higher glycolytic rates compared to the non-regenerating species (ANOVA, *F*=23.3301, *p*< 0.0001) (d-e). Also, *Acomys* and *Oryctolagus* fibroblasts exhibited significantly higher glucose sensitivity compared to *Mus* and *Rattus*, responding rapidly to glucose injection 10 to 15 minutes faster (d). Individual data points represent biological replicates for each species (n=4). FCCP: carbonyl cyanide 4-(trifluoromethoxy)phenylhydrazone. 2-DG: 2-Deoxy-d-glucose.

In parallel with the MST, we ran a glycolytic stress test measuring extracellular acidification rates (ECAR) while sequentially injecting glucose, oligomycin and 2-deoxyglucose to calculate glycolytic rate, glycolytic capacity, glycolytic reserve and non-glycolytic acidification (Fig. 1d and Supp. Fig. S1d-f). Measuring the amount of extracellular acid produced by cells from all four species we observed significantly higher glycolytic rates in cells from the regenerating species (Fig. 1d-e). Although we did not detect significant differences among other GST parameters between regenerators and non-regenerators, we observed that *Acomys* and *Oryctolagus* showed a similarly high sensitivity to glucose, responding immediately to extracellular glucose while *Mus* and *Rattus* exhibited a delayed reaction (Fig. 1d). Glycolytic reserve and percent glycolytic reserve levels were lowest in *Acomys* among the species assessed (Supp. Fig. S1d, f), which was interesting considering their high glucose sensitivity and glycolytic rate (Fig. 1 d-e). Taken together, these results show that *Acomys* and *Oryctolagus* fibroblasts exhibit a low basal metabolic rate and high basal glycolytic rate, while *Mus* and *Rattus* fibroblasts exhibit high maximal respiration and low glucose sensitivity. *Oryctolagus* fibroblasts exhibit an increased sensitivity to glucose but also an ability to ramp up mitochondrial respiration abruptly in response to an acute energy deficit.

To extend our measures of cellular metabolism, we next ran untargeted and targeted metabolomics on primary fibroblasts from *Acomys* and *Mus*. Initial untargeted analysis with hundreds of recovered compounds recovered a strong association by species (Supp. Fig. S2a-d and Supp. Tables 1a-b). Armed with this validation, we next targeted key glycolytic and TCA cycle intermediates. We observed significantly higher levels of glycolytic intermediates, namely glucose-6-phosphate, dihydroxyacetone phosphate/glyceraldehyde-3-phosphate (DHAP-G3P) and lactic acid (Fig. 2a). We also observed increased levels of fumaric acid, oxaloacetate (OAA) and alpha-ketoglutaric acid (αKG) in *Acomys* fibroblasts. Oxidized and reduced forms of glutathione were also highly elevated in *Acomys* compared to *Mus* (Fig. 2a), supporting previous work showing that *Acomys* fibroblasts exhibit increased glutathione peroxidase activity in response to induced oxidative stress (18). We also observed higher levels of the reduced and oxidized forms of the coenzyme NADP in *Acomys*. Together, our metabolomics screen reinforced a reliance on glycolysis in *Acomys* fibroblasts and provide additional support for increased antioxidant activity associated with a large, reduced glutathione pool.

**Figure 2.**
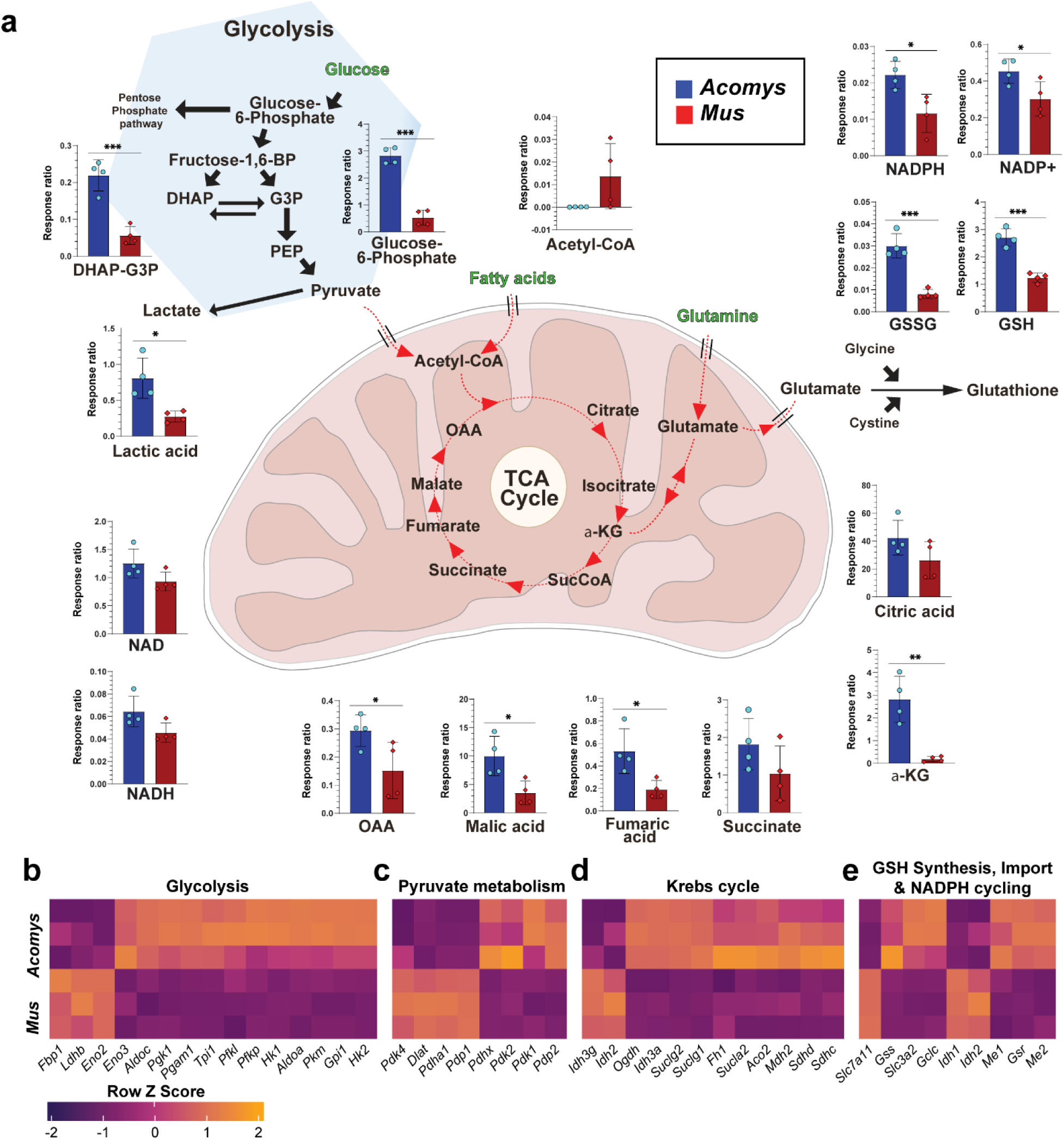
Metabolomics and transcriptomic comparison of *Acomys* and *Mus* fibroblasts. Untargeted metabolomics indicates *Mus* and *Acomys* fibroblasts (4 biological replicates/species) group distinctly. (**a**) Targeted metabolomics analysis shows differential metabolite levels for glycolysis, the Krebs cycle and glutathione synthesis pathways. *Acomys* fibroblasts have significantly higher levels of glucose-6-phosphate, lactic acid, dihydroxyacetone phosphate as well as high glutathione levels. Levels of TCA intermediates AKG, malic, fumaric and oxaolacetic acid were significantly higher in *Acomys*. Student’s t-tests were used for species-species comparisons of individual metabolite levels. (* *p*<0.05, ** *p*<0.001, *** *p*<0.0001). (**b-e**) Heatmaps generated from bulk RNA-seq comparison of *Acomys* and *Mus* fibroblasts (n=3/species) showing per-gene normalized expression of key enzymes involved in glycolysis (b), pyruvate metabolism (c), TCA cycle (d) and glutathione synthesis & recycling (e). All genes included in the heatmap are significantly-differentially expressed across biological replicates (n=3) between the two species (FDR<0.05).

After observing key metabolite differences between *Acomys* and *Mus* fibroblasts, we generated bulk RNA-seq transcriptomes from cultured fibroblasts. To assess metabolic genes, we used the MitoCarta3.0 database (33) to derive gene sets involved in oxidative phosphorylation, ROS management and glucose metabolism (i.e., glycolysis, pyruvate metabolism, TCA cycle, and glutathione synthesis and recycling). This analysis revealed that the glycolytic enzymes aldolase c (*Aldoc*), triose phosphate isomerase (*Tpi1*), hexokinases 1 & 2 (*Hk1& Hk2*), phosphoglyceromutase, and phosphoglycerate kinase (*Pgk1*) were highly elevated in *Acomys* compared to *Mus* (Fig. 2b). Enolase exhibits a species-specific expression pattern, with *Eno3* significantly higher in *Acomys* and *Eno2* higher in *Mus.* Key components of the pyruvate dehydrogenase complex (*Pdh*), which catalyzes the conversion of pyruvate to acetyl coA, namely dihydrolipoamide S-acetyltransferase (*Dlat*) and pyruvate dehydrogenase E1 subunit alpha 1 (*Pdha1*), were highly expressed in *Mus* compared to *Acomys*. Pyruvate dehydrogenase kinases (*Pdk1* & *Pdk2*), which inhibit pyruvate conversion to acetyl coA by inactivating pyruvate dehydrogenase via phosphorylation, were elevated in *Acomys* (Fig. 2c). The blocking of *Pdh* and the concurrent upregulation of *Ldha* in *Acomys* (Suppl. Table 2) support that glucose consumed is being shunted towards lactate production instead of being fed into the TCA cycle. Lactate dehydrogenase B (*Ldhb*), which catalyzes the conversion of lactate back to pyruvate, was also elevated in *Mus* (Fig. 2b), corroborating the low levels of basal ECAR observed earlier (Fig. 1d-e). The upregulation of pyruvate dehydrogenase phosphatase catalytic subunit 1 (*Pdp1*) which activates inactivated Pdh via dephosphorylation was also elevated in *Mus*, further supporting the production of acetyl coA from cytosolic pyruvate (Fig. 2a). Interestingly, there were more TCA cycle enzymes upregulated in *Acomys* (Fig. 2d), mirroring the higher TCA intermediates observed in the mass spectrometry runs (Fig. 2a), further pointing to an unusual increase in TCA cycle flux without the expected oxygen consumption rates (Fig. 1a-c). In *Acomys*, we also observed elevated expression of glutathione synthetase (*Gss*), glutamate-cysteine ligase catalytic subunit (*Gclc*), and glutathione reductase (*Gsr*), the critical enzymes in the synthesis and recycling of glutathione compared to *Mus* (Fig. 2e), supporting the high levels of both reduced and oxidized glutathione observed earlier (Fig. 2a). Malic enzyme 1 (*Me1*) and malic enzyme 2 (*Me2*), which generate NADPH downstream of the export of TCA intermediates into the cytosol, were also highly expressed in *Acomys*, in part supporting the high levels of reduced glutathione observed in the targeted metabolomics run (Fig. 2a).

### Functional mitochondria in Acomys fibroblasts present as large spheres

Next, we asked if mitochondria in regenerating species exhibited morphological differences that could explain their observed predilection for glycolysis using CMX-Ros Mitotracker red (MR) and Mitotracker green (MG). MR labels mitochondria with a high resting membrane potential which are often interpreted as active. *Mus* mitochondria stained with MR exhibited the classical morphology typified by tubular networks extending throughout the cytoplasm (Fig. 3a-a’). In stark contrast, *Acomys* fibroblasts presented large, spheroid mitochondria clustered around the nucleus which were relatively few in number (∼20-40/cell) and resembled previously reported megamitochondria with respect to shape (34) (Fig. 3c-c’). *Oryctolagus* and *Rattus* fibroblasts exhibited tubular mitochondrial networks more in line with *Mus*, although a range of small to large spherical mitochondria were also observed randomly dispersed in their cytoplasm (Fig. 3b-b’, d-d’). The circular mitochondria observed in *Oryctolagus* and *Rattus*, however, were smaller, most approximating the size of fragmented mitochondrial tubes in contrast to the large mitochondria found in *Acomys* (Fig. 3c-c’).

**Figure 3.**
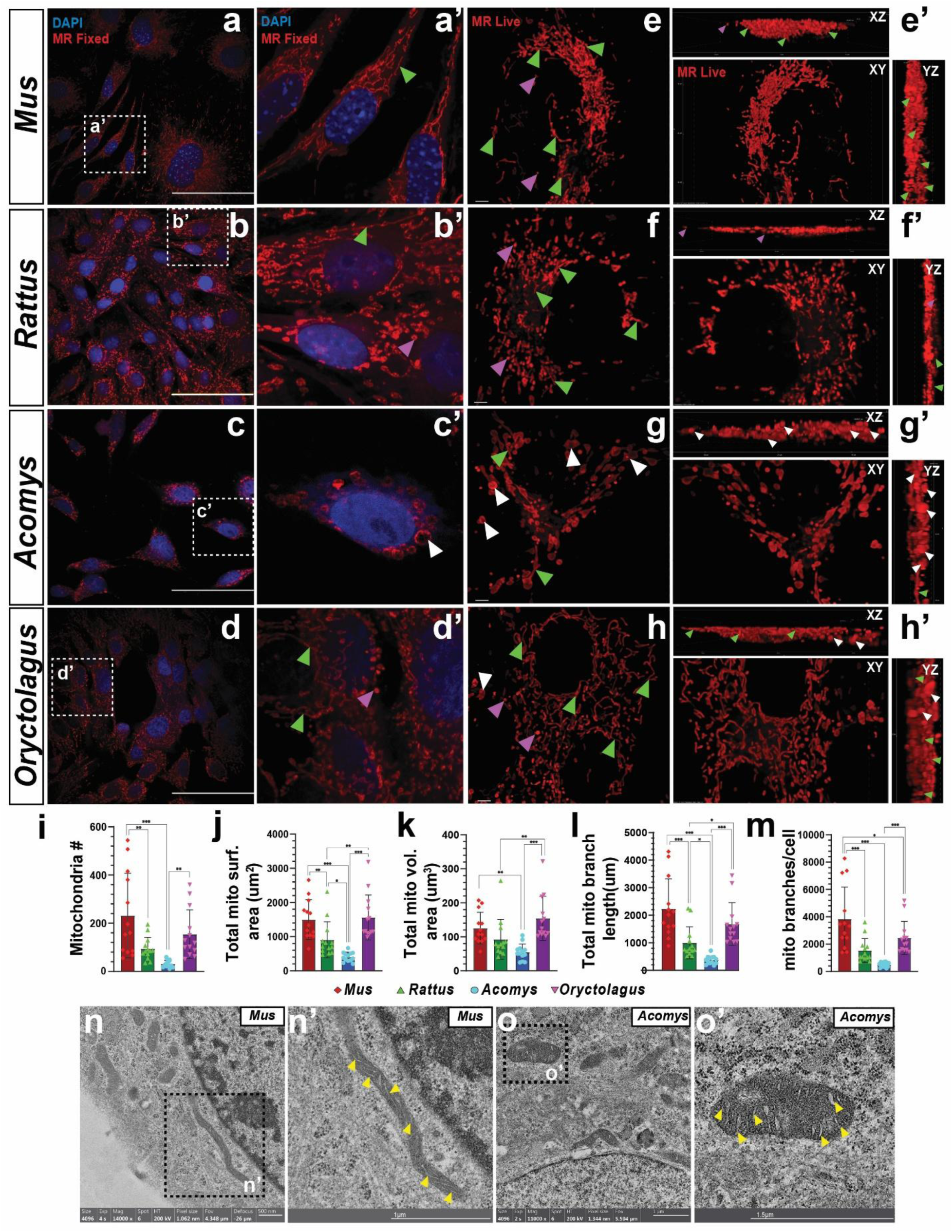
Active mitochondria in healthy *Acomys* cells are large and spherical. (a’-d’) Representative images of fixed fibroblasts from *Mus, Acomys, Rattus* and *Oryctolagus* stained with Mitotracker CMXROS red dye (MR). (**a-d**) Inset images at 60x magnification show classic tubular mitochondria in *Mus*, *Rattus* and *Oryctolagus* (green arrows), whereas active mitochondria in *Acomys* appear as large, spheres (white arrows) (Scale bars: 80 μm). Small, round mitochondria were also visible in *Rattus* and *Oryctolagus* (purple arrows). (**e-h**) High resolution, maximal projection images of live fibroblasts stained with MR and imaged at 4 x 60X. (Scale bars: 2μm). (**e’-h’**) Representative live images of cultured P2 ear pinna fibroblasts from *Mus, Acomys, Rattus* and *Oryctolagus* stained with MR showing of cells from (e-h) projections showing xz, and yz, planes to indicate mitochondrial shape in 3D (Scale bars: 2 μm). Green arrows and purple arrows point to tubular and small spherical mitochondria respectively. White arrows point to massive spheroid mitochondria. (**i-m**) Mitochondrial analysis of 3D images in (e-h) indicate differences in mitochondrial count per cell (ANOVA, *F*= 8.8026, *p*<0.0001), total mitochondrial surface area (ANOVA, *F*= 14.5368, *p*<0.0001), total mitochondrial volume (ANOVA, *F*= 9.0002, *p*<0.0001), mitochondrial branch length (ANOVA, *F*= 15.8517, *p*<0.0001) and number of mitochondrial branches per cell (ANOVA, *F*= 13.5591, *p*<0.0001) in the four species. Post-hoc pairwise comparisons were carried out using the Student’s *t*-test. (* *p*<0.05, ** *p*<0.001, *** *p*<0.0001). (**n-o**) TEM images of fibroblasts obtained from young *Mus* and *Acomys*. Yellow arrows indicate intact cristae.

To determine if the mitochondria phenotype observed in *Acomys* was strictly associated with active mitochondria, we used MG to label mitochondrial protein cysteine residues that accumulate in the mitochondrial matrices independent of membrane potential to label all mitochondria (35, 36). In *Mus*, *Rattus* and *Oryctolagus*, the mitochondrial networks observed with MG mirrored that observed with MR (Supp. Fig. S3a, c-d). While MG staining revealed classic tubular networks in *Acomys*, it did not stain large, spherical mitochondria (Suppl. Fig. S3b). Because the chemistry of MitoTracker green staining appeared to interfere with the ability to stain all *Acomys* mitochondria, we examined live cells stained with MR using 3-dimensional high-resolution imaging and observed that *Mus* fibroblasts maintained extensive mitochondrial tubular networks with several small punctate mitochondria resembling fission products (Fig. 3e-e’). *Rattus* and *Oryctolagus* fibroblasts also showed extensive tubular networks with additional small, spherical mitochondria interspersed in the cytoplasm (Fig. 2f-f’, h-h’). In *Acomys,* we observed some tubular mitochondria (Supp. Fig. S3e), but the dominant morphology proved to be massive, spherical mitochondria, with some also yielding an oblong or tubular shape (Fig. 3g-g’ and Supp. Fig. S3f).

Most of the mitochondria in *Acomys* appeared distinctly larger than those seen in the other three species and *Acomys* mitochondria exhibited distinctly lower density. Maximum projection through 3-dimensional stacks revealed that the massive mitochondria observed in *Acomys* were spherical, not cylindrical in nature (Fig. 3g-g’). The tubular mitochondria observed in the other species formed extensive networks (Fig. 3e-h and Supp. Videos 1-4). Quantifying 3D mitochondrial morphology using the Mitochondrial Analyzer plugin on ImageJ (37), showed that *Acomys* fibroblasts had significantly fewer mitochondria compared to *Mus* and *Oryctolagus* (Fig. 3i). *Acomys* mitochondrial surface area was significantly lower amongst the four species (Fig. 3j), and total mitochondrial volume was significantly smaller compared to *Mus* and *Oryctolagus* (Fig. 3k). *Acomys* had fewer mitochondrial network branches (Fig. 3m) and shorter branches per cell (Fig. 3l). These patterns in *Acomys* were consistent across the number of branch junctions per cell/mitochondrial volume (Supp. Fig. S3i-j), as well as branch endpoints per cell/mitochondrial volume (Supp. Fig. S3k-l), supporting few tubular mitochondrial networks.

Since large, spherical mitochondria in mice and humans have been linked to poor respiratory measures due to underdeveloped cristae formation (38), we used TEM imaging to examine the cristae structure in mitochondria from *Mus* and *Acomys* (Fig. 3n-o’ and Supp. Fig. S3m-n’). While mostly tubular mitochondria with intact cristae were observed in *Mus*, TEM images of large, spherical mitochondria in *Acomys* also revealed well-developed, intact cristae supporting their activity (Fig. 3n-o’ and Supp. Fig. S3m-n’). Considered together, these results demonstrate that while *Mus*, *Rattus* and *Oryctolagus* possess extensive networks of mostly active, tubular mitochondria, *Acomys* possesses mostly large, spherical mitochondria with well-developed cristae distinguishing them from megamitochondria found in pathological mouse and human cells (34).

Having generated expression data for *Acomys* and *Mus* fibroblasts, we asked whether differential expression of critical mitochondrial fusion and fission genes might correlate with the observed mitochondrial phenotypes. We used the MitoCarta3.0 database to derive gene sets regulating mitochondrial shape dynamics (Supp. Fig. S4a-c). Some genes facilitating mitochondrial fission were upregulated in *Mus* (i.e., *Fis1*, *Armc10*, and *Mtfr1*), while others were upregulated in *Acomys* (i.e., *Mief1*, *Mief2, Mtfp1, Mtfr2, Mff, Mul1*) (Supp. Fig. S4a). Similarly, some genes regulating mitochondrial fusion were significantly elevated in *Acomys* (i.e., *Mfn1*), but others were significantly elevated in *Mus* (i.e., *Mfn2, Miga1, Arl2*) (Supp. Fig. S4b). Moreover, core fusion (*Opa1*) and fission (*Dnm1* - coding for *Drp1*) genes were similarly expressed in both species (Supp. Fig. S4c). Together, these data do not point towards inherent species-specific expression levels of fusion and fission genes as correlates with observed differences in mitochondrial morphology.

### Acomys mitochondria are inherently depolarized but appear highly efficient at producing ATP

Our morphological investigation suggested membrane polarization as a source of variation across species that might explain the unique metabolic state observed in spiny mouse cells. We turned to JC-1 to visualize and quantify mitochondrial function. First, we used microscopy to visualize JC-1 green and red fluorescence from all four species which revealed reduced mitochondrial density around the nucleus in *Acomys* compared to the other three species (Fig. 4a-b). Curiously, we noticed that only a fraction of *Acomys* cells showed red aggregations from the JC-1 stain (Fig. 4a). To estimate mitochondrial function using JC-1, we used a plate reader to quantify red and green emission intensities (Fig. 4c). Culturing cells to ∼80% confluency, we incubated with JC-1 for 30 minutes, washed and then simultaneously detected emissions at 529 nm and 590 nm (Fig. 4d-f’). Using this method, membrane potential (red to green signal ratio) in *Acomys* fibroblasts was significantly lower compared to the other three species (Fig. 4d-d’). We also exposed biological replicates to two additional treatments: oligomycin and FCCP, in a bid to induce mitochondrial hyperpolarization and depolarization respectively (Fig. 4e-f’). In response to oligomycin, *Acomys* mitochondria exhibited a two-fold increase in membrane potential as expected since inhibiting ATP synthase produces a net positive build-up of protons in the mitochondrial intermembrane space resulting in mitochondrial hyperpolarization (Fig. 4e-e’). Interestingly, *Rattus*, *Mus*, and *Oryctolagus* fibroblasts showed no significant change in mitochondrial membrane potential in response to oligomycin treatment suggesting that their mitochondria, on average, were hyperpolarized in culture (Fig. 4e-e’). The inverse was seen for the FCCP treatment, where proton gradient disruption triggered dramatic depolarization in *Mus*, *Rattus* and *Oryctolagus* mitochondria as expected, however *Acomys* fibroblasts showed no significant decrease in membrane potential (Fig. 4f-f’). Thus, our JC-1 data further support the assertion that *Acomys* mitochondria are generally low-functioning, and yet they are very sensitive to ATP synthase inhibition, suggesting they are highly efficient with respect to ATP production.

**Figure 4.**
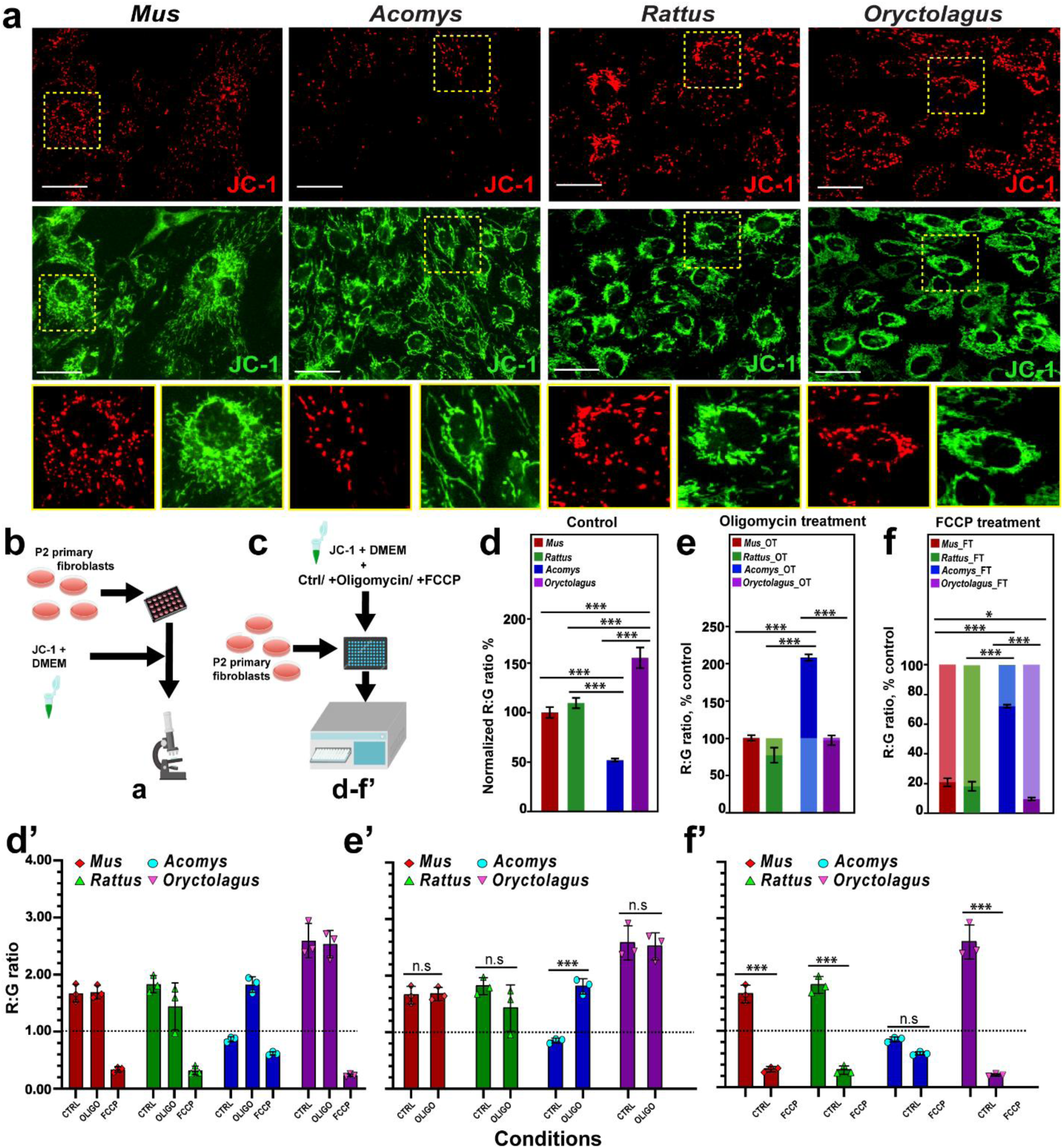
*Acomys* mitochondria have exhibit low membrane potential and are tightly coupled. **(a)** Live fluorescence images of P2 ear pinna fibroblasts from *Mus*, *Acomys*, *Rattus* and *Oryctolagus* stained with JC-1 and imaged in red and green channels at 20x magnification, (Scale bars = 20 µm). **(b-c)** Schematics indicating how samples were observed initially via fluorescence microscopy and subsequently using a faster plate scan. **(d-f’)** Due to the photolability of the JC-1 dye, fluorescence intensity was better measured using a plate reader to capture acute changes in membrane potential responding to mitochondrial inhibitors FCCP and Oligomycin. Red:green signal ratios were quantified and normalized to total µg protein and pegged against untreated *Mus* (**d-f**). *Acomys* fibroblasts exhibited the lowest membrane potential among all four species (51.9% of *Mus*), with *Oryctolagus* having the highest (155.4% of *Mus*), with *Rattus* (109%) not significantly different from *Mus* (ANOVA, *F*= 167.4372, *p*<0.0001) (**d-d’**). Membrane potential in *Acomys* mitochondria increased two-fold in response to oligomycin, while in *Rattus*, *Mus*, and *Oryctolagus* exhibited no significant change compared to basal membrane potential (ANOVA, *F*= 10.0687, *p*=0.0006) (**e, e’**). FCCP treatment caused an insignificant decrease in mitochondrial membrane potential in *Acomys*, compared to *Mus*, *Oryctolagus* and *Rattus* (*p*_Mus_<0.001, *p*_Oryctolgagus_<0.001, *p*_Rattus_<0.001,) which dropped significantly to 20.4%, 9.2%, and 17.8% respectively of their basal red:green ratios in response to the uncoupler (ANOVA, *F*= 60.1244, *p*<0.0001) (**f, f’**). Individual data points represent biological replicates per species (n = 3). Post-hoc pairwise comparisons were carried out using the Student’s *t*-test. (* *p*<0.05, ** *p*<0.001, *** *p*<0.0001).

### Frozen, isolated mitochondria from Acomys and Oryctolagus share low Complex I, II, and IV dependent respiration

After observing key differences in mitochondrial membrane potential across all four species, we set out to quantify oxygen consumption from isolated mitochondria. Isolated mitochondria were frozen and assayed using the electron transport chain protocol (ETCP) (39), where we utilized alamethicin and supplemented lost cytochrome c. This method allows assessment of mitochondrial complex function without a contribution from the proton gradient (40), which is disrupted after successive freeze-thawing cycles (41). Despite losing the proton-motive force, activity from complexes I, II and IV are observed, allowing the assessment of ETC complex activity in archived samples, years after collection (39, 40). Performing the ETCP assay we observed *Oryctolagus* mitochondria to possess significantly reduced complex I, II and IV activities (oxygen consumption per microgram of mitochondrial protein) compared to *Mus* and *Rattus* (Fig. 5a-e). Similarly, *Acomys* activity in complexes I, II and IV was significantly lower compared to the non-regenerators (Fig. 5a-e). When complex activity is considered in toto across the entire treatment time course, a stark dichotomy emerges separating the regenerators and non-regenerators; *Acomys* and *Oryctolagus* mitochondria show consistently lower oxygen consumption per µg of mitochondrial protein even in response to TMPD and Ascorbic acid, which restored oxygen consumption to levels observed prior to Antimycin A injection (Fig. 5a). Taken together, in the absence of a protonmotive gradient, mitochondria from regenerators exhibit intrinsically lower oxygen consumption compared to non-regenerators.

**Figure 5.**
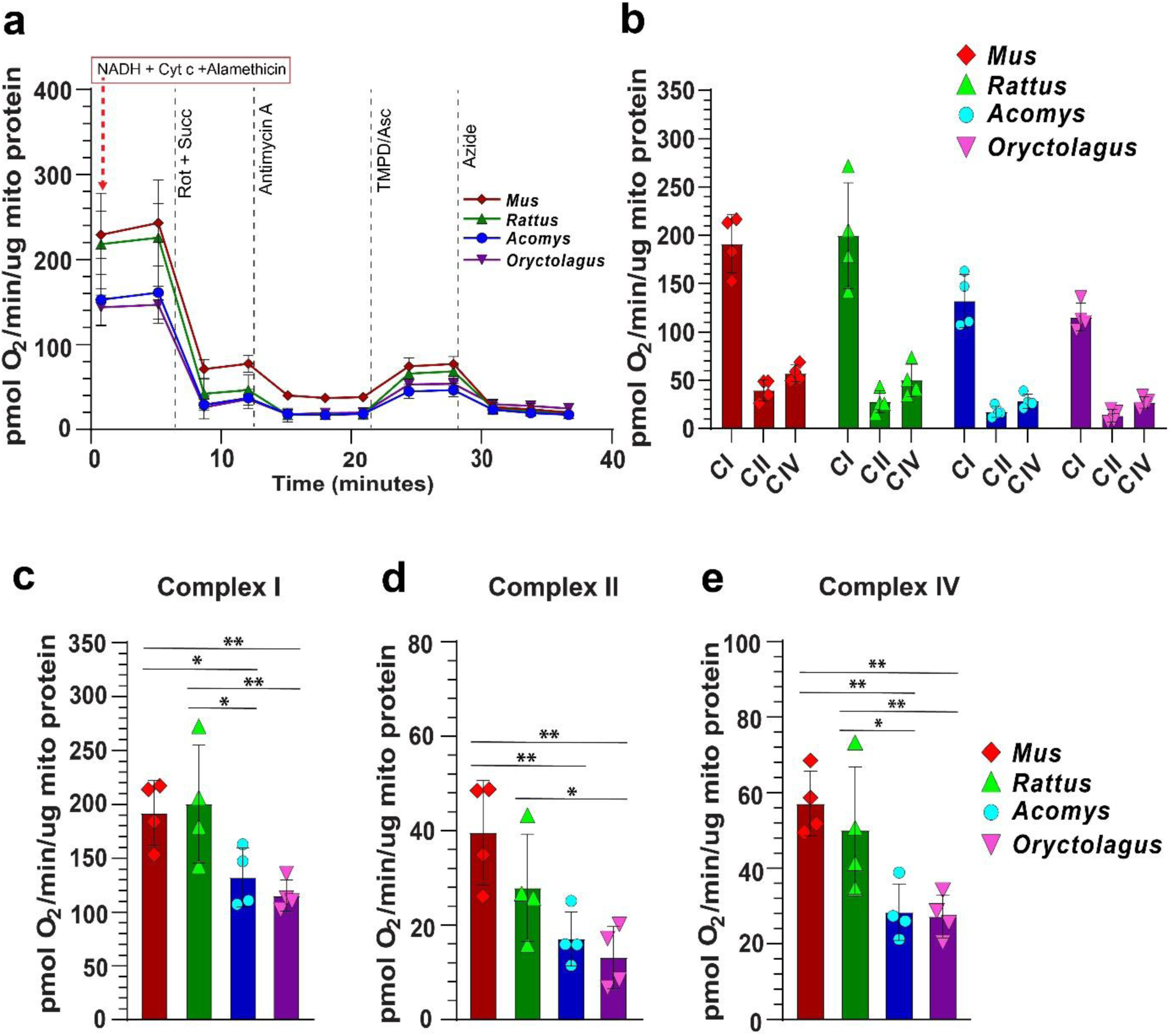
*Mus* and *Rattus* fibroblasts have significantly elevated ETC complex activity compared to *Acomys* and *Oryctolagus*. **(a-e)** Representative seahorse traces of oxygen consumption per microgram of mitochondrial protein quantified in isolated, frozen mitochondria from P2 ear pinna fibroblasts using ETCP, a modified version of the RIFS respirometry assay. Frozen mitochondria were assayed utilizing alamethicin, NADH and supplementing lost cytochrome c. *Oryctolagus* and *Acomys* mitochondria had lower (CI) Complex I-dependent respiration compared to *Mus* and *Rattus* (ANOVA, *F*=5.8468, *p*=0.0106). Similarly, (CII) Complex II-dependent respiration in *Oryctolagus* was again lower than in the non-regenerators. *Mus* Complex II respiration was higher than in *Acomys*, although the difference between *Acomys* and *Rattus* was not significant (ANOVA, *F*=6.8119, *p*=0.0062). Cytochrome c oxidase (CIV) respiration was significantly higher in the non-regenerators than the two regenerators (ANOVA, *F*=8.3265, *p*=0.0029). For all three Complex-dependent respiration measurements, individual data points represent biological replicates/species, (n = 4). An average of four technical replicates were used to represent each biological replicate. Post-hoc pairwise comparisons were carried out using the Student’s *t*-test. (* *p*<0.05, ** *p*<0.001, *** *p*<0.0001).

### Large, spherical mitochondria persist across lifespan and remain resilient to oxidative stress in spiny mice fibroblasts

To further assess how cellular metabolic state and mitochondrial morphology might functionally affect fibroblast phenotype we examined *Mus* and *Acomys* cells isolated from embryonic (*Mus* - E12.5, *Acomys* - E20-25), young (*Mus* - 3 months, *Acomys* - 6 months) and old (*Mus* - 2+ years, *Acomys* - 4+ years) animals (Fig. 6a-f). Staining *Mus* cells with Mitotracker red, we observed a network of small, round and tubular mitochondria in E12.5 MEFs that transitioned into the largely tubular network associated with cells from adult animals (young or old) (Fig. 6a). In stark contrast, MR staining of *Acomys* fibroblasts across lifespan revealed that large, spherical mitochondria were the predominant, active form around the nucleus and in the cytoplasm independent of age (Fig. 6a). Subsequent cellular TEM analysis across all ages reinforced the retention of large, spherical mitochondria in *Acomys* fibroblasts (Fig. 6b). These data support that the unique mitochondrial phenotype found in healthy *Acomys* fibroblasts is independent of development and aging supporting evolved functionality related to cellular physiology.

**Figure 6.**
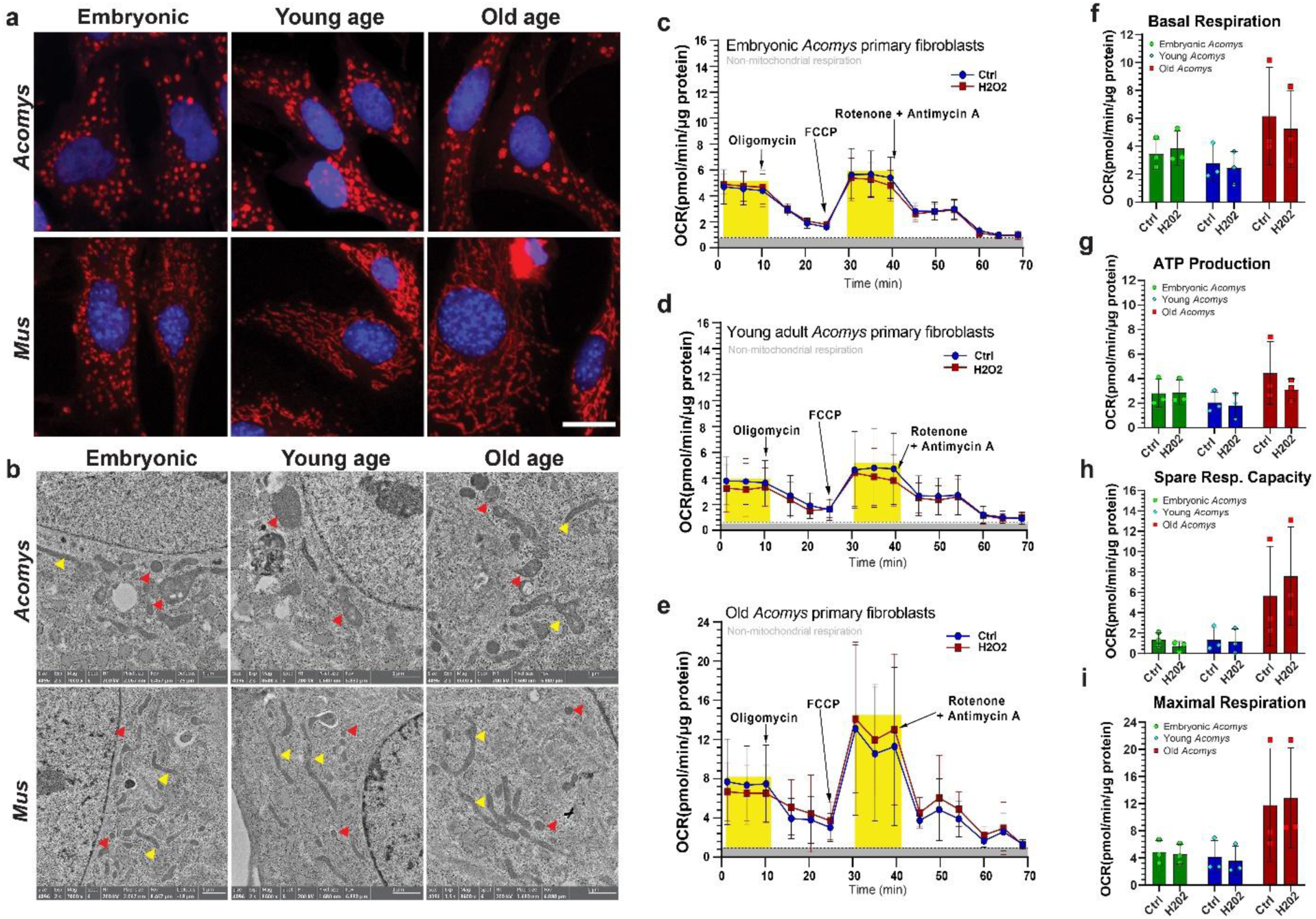
*Acomys* primary fibroblasts maintain large, spherical mitochondria and are highly resistant to oxidative stress across lifespan. (a) Primary ear pinna fibroblasts from fetal, young and old *Mus* and *Acomys* stained with MitoTracker Red (scale bar = 20 µM). *Acomys* fibroblasts exhibit mostly large, spherical mitochondria across all ages. Small, circular mitochondria are also present in *Mus* embryonic fibroblasts, but mitochondria adopt a tubular morphology after birth. (b) Electron micrograph images showing spherical and tubular mitochondria in *Acomys* and *Mus* (red arrows = round mitochondria, yellow arrows = tubular mitochondria). (c-i) Mitochondrial function in *Acomys* is unaffected by 300 µM H_2_O_2_ treatment across different age classes as measured by OCR (oxygen consumption rates) via the Seahorse mitochondrial stress test. There aren’t significant changes in Basal respiration (ANOVA, *F*=0.1576, *p* = 0.8559), ATP production (ANOVA, *F*= 0.4455, *p*= 0.6507), Maximal respiration (ANOVA, *F*= 0.0449, *p* = 0.9563), and Spare respiration capacity (ANOVA, *F*= 0.3455, *p*= 0.7147), in response to H_2_O_2_ treatment. Individual data points represent biological replicates (n = 3) per age class/species. Post-hoc pairwise comparisons were carried out using the Student’s *t*-test. (* *p*<0.05). FCCP: Carbonyl cyanide-p-trifluoromethoxy phenyl hydrazone.

Previous work in our lab uncovered that fibroblasts from young spiny mice were resilient to very high levels of hydrogen peroxide (18) and thus we exposed cells from all ages to 300µm H_2_O_2_ and assessed mitochondrial stress (Fig. 6c-i). Two trends were noted. First, cells from the oldest animals exhibited a slight, but insignificant increase in basal respiration (Fig. 6f), spare respiratory capacity and maximal respiration (Fig. 6h-i). Despite these trends, when cells from all age groups were exposed to 300µm H_2_O_2_ there was no significant alteration in metabolic function underscoring the resiliency of spiny mouse cells to oxidative stress (Fig. 6c-i). Together, our data supports that a unique mitochondrial phenotype, low metabolic rate, and high resilience to oxidative stress are features of spiny mouse cells that are invariant across lifespan. The degree to which these cellular phenotypes functionally support regeneration awaits further testing.

## DISCUSSION

This study set out to characterize cellular metabolism in primary fibroblasts from two highly regenerative mammals (*Acomys* - spiny mice and *Oryctolagus* - rabbits) compared to fibroblasts from non-regenerative *Mus* and *Rattus*. By quantifying cellular oxygen consumption (OCR) and extracellular acidification (ECAR) in primary ear pinna fibroblasts we found that *Acomys* and *Oryctolagus* cells preferentially used glycolysis to support basal metabolism in culture and exhibited increased sensitivity to glucose compared to poorly regenerative *Mus* and *Rattus*. Conversely, *Mus* fibroblasts showed high basal respiration, maximal respiration, and spare respiratory capacity. We also observed high levels of glycolytic intermediates and glutathione (oxidized and reduced) in *Acomys*, as well as some TCA intermediates. Interesting among them was the stark elevation of αKG in *Acomys*, despite unremarkable levels of the upstream TCA intermediate, citric acid. Apart from isocitrate oxidation, αKG can be generated from the systematic deamination and oxidation of glutamine and glutamate respectively. αKG can also be transaminated to form glutamate which can feed into glutathione synthesis. We observed high levels of both glutathione and αKG in *Acomys*, pointing to a possible rerouting of TCA cycle intermediates via αKG transamination towards increased glutathione synthesis. Utilizing radiolabeled glucose or glutamine for metabolite tracing in future studies will provide a potent test of this hypothesis. The combined phenotype of high glycolytic flux, elevated TCA cycle with concurrent low oxygen consumption suggests that the TCA cycle in *Acomys* is decoupled from the electron transport chain. Still, our physiological and metabolomic data clearly point towards an inherent glycolytic bias in *Acomys* ear pinna fibroblasts and those from *Oryctolagus* as well.

This study also revealed a unique mitochondrial phenotype in *Acomys* fibroblasts distinct from the well-established tubular, networked morphology typically seen in fibroblasts from adult mammals, including humans. Specifically, *Acomys* mitochondria resembled megamitochondria (34, 42), albeit functional, presenting as large, active spheroids, whereas mitochondria in *Mus* fibroblasts were almost completely tubular, exhibiting extensive networks dispersed throughout the cytoplasm. *Rattus* and *Oryctolagus* were similar to *Mus* but also exhibited some small spherical mitochondria. High resolution 3-dimensional renderings of live fibroblasts from all four species stained with Mitotracker CMXRos red stain supported the tubular phenotypes in *Mus*, *Rattus* and *Oryctolagus*. In contrast, *Acomys* showed a range of mitochondrial phenotypes dominated by large spheroid phenotypes but also revealed randomly scattered tubes. Independent of morphology, mitochondrial density was consistently lower in *Acomys* compared to the other three species. Although we are as of yet unable to explain why mitotracker green does not consistently stain the large, spherical mitochondria in *Acomys*, recent studies have shown that its staining success is affected by membrane potential (43) as well as damage or dysfunction due to oxidative stress (44), fission/fusion events and autophagy.

Low respiring, small, circular mitochondria are usually associated with embryonic stem cells (25) where they are ascribed poor respiratory measures due to underdeveloped cristae formation and an anoxic environment (26). Networks of tubular mitochondria create increased surface area for greater oxidative phosphorylation and are thus associated with high oxygen consumption. To explore how form might echo function in primary fibroblasts, we employed a JC-1 assay to measure intracellular mitochondrial membrane potential. To understand how the JC-1 assay helps us quantify electron transport chain (ETC) function, consider the following analogy. Utilizing FCCP (membrane depolarizer) and oligomycin (proton channel blocker) allows us to conceptualize the entire ETC as a tap-connected garden hose with an operable spout. Turning on the tap represents complex I & II reactions, with electron transfer from NADH (and FADH_2_) to ubiquinone simultaneously pumping protons from the mitochondrial matrix into the intermembrane space. The differential membrane potential drives protons back into the matrix via ATP synthase, which couples ATP production to proton pumping back into the matrix as water flows through the hose. Administering oligomycin stops up the garden hose at the outflow thus generating a significant increase in mitochondrial membrane potential due to the buildup of protons in the intermembrane space compared to the matrix. Adding FCCP is analogous to punching holes in our hose which decreases net pressure of water flowing through the hose causing membrane potential to fall. After oligomycin treatment, a spike in membrane potential was only observed in *Acomys*, suggesting that the mitochondria in the other species were maximally polarized most likely due to a higher respiratory rate. When FCCP was added, we observed a significant drop in mitochondrial membrane potential in *Mus*, *Rattus* and *Oryctolagus*. Surprisingly, adding FCCP did not decrease membrane potential in *Acomys* suggesting that *Acomys* mitochondria are intrinsically depolarized. Together, these data support that *Acomys* mitochondria are in fact tightly coupled and highly efficient resembling a metabolic state sometimes observed in ESCs (45).

To further dissect function and efficiency of the ETC in these cells, we utilized the Electron Transport Chain Protocol assay to measure intrinsic mitochondrial oxygen consumption and ATP production of individual complexes in the absence of a proton gradient (39, 40). Deploying this assay, we observed a trend that split mitochondria from the four species into two groups: regenerating and non-regenerating exhibiting low and high oxygen consumption respectively. *Mus* and *Rattus* showed consistently higher levels of oxygen consuming capacity with high activity observed for complexes I, II and IV. Importantly, while our JC-1 data demonstrated that *Oryctolagus* mitochondria appeared similar to *Mus* and *Rattus* with respect to membrane potential, data from our ETCP analysis provides a potential explanation for this discrepancy. Here we observed decoupling between high membrane potential observed in rabbit fibroblasts and actual oxygen consumption capacity per microgram of mitochondrial protein. Although mitochondria from rabbit ear pinna fibroblasts exhibited high membrane potential, this does not reflect in their low basal respiration levels and high glycolytic flux mirroring that of *Acomys*. The high mitochondrial membrane potential observed in *Oryctolagus* interestingly supports their high spare respiratory capacity, which *Acomys* do not share. Furthermore, our data suggests that low basal respiration and maximal respiration levels (forced by FCCP) observed in *Acomys* are not just the result of low mitochondrial numbers, but instead, these mitochondria are depolarized and intrinsically inefficient (per µg of mitochondrial protein) at consuming oxygen explaining their low oxidative phosphorylation levels. Further studies will be necessary to determine a deeper mechanistic explanation for the phenotypes observed in *Oryctolagus* i.e., the discrepancy between high glycolytic rates, the low basal respiration levels, and intrinsic inefficiency in oxygen consumption despite having higher mitochondrial membrane potential.

Considering the influence of aging on metabolic phenotype, mitochondrial labelling and TEM imaging carried out indicated that *Acomys* fibroblasts maintain their unique mitochondrial phenotype across lifespan and actively resist senescence inducing stressors. A key finding here is that fibroblasts from fetal and old *Acomys* concurrently resisted ROS-induced mitochondrial dysfunction as earlier observed young adult fibroblasts (18). Therefore, this concurrent persistence of cellular resilience and metabolic phenotype gives more credence to the hypothesis that this metabolic signature is a strong causal factor. Also, it could be argued that this mitochondrial phenotype in *Acomys* is induced by hypoxic culture conditions (3% O_2_). It is key to note that this oxygen concentration better represents physiological levels in vivo, and that cells from the other species observed were cultured together in the same conditions but did not share this mitochondrial phenotype and that this mitochondrial state in *Acomys* may be a species-specific adaptation.

Research performed over the last decade, primarily in laboratory mice, identified that certain pre-injury fibroblast phenotypes appeared to support regenerative healing over fibrotic repair (46–50). While some of these studies connected phenotype to specific signaling pathways (e.g., Wnt-signaling activation during regeneration) (46, 51), others uncovered differential mechano-sensing (52, 53) or inflammatory priming (54) as key drivers of skin regeneration. Although deploying these approaches has yet to successfully alter healing of more complex tissue injuries, such as large (>4mm) ear pinna defects or amputated P2 digit tips, the general principle that specific fibroblast phenotypes are at least partially required for complex tissue regeneration is well-supported from contemporary regeneration studies (31, 55, 56). Identifying molecular regulators of fibroblast phenotype beyond transcriptomic profiling remains poorly characterized, but specific metabolic characteristics are emerging as key drivers underlying cellular phenotypes. Although it can be argued that this pro-glycolytic state observed in *Acomys* and *Oryctolagus* may largely be responsible for a potential towards increased cell proliferation in the wound healing context, a previous study indicated that non-regenerative, OxPhos-dependent *Rattus* fibroblasts, although susceptible to ROS-induced senescence, were highly proliferative even more than observed in *Acomys* and the other non-regenerator, *Mus* (18). It is important to point out that the data collected from *Acomys* fibroblasts (high glycolytic and TCA flux (57, 58), with concurrent low mitochondrial respiration) in this study, suggest that beyond having a glycolytic bias, these fibroblasts seem to exhibit aspects of the Warburg program described in cancer cells. The elevated expression of master regulators of this program, namely *Hif1α*, *Myc* and *Akt1* & *2* in *Acomys* as well as some of their downstream targets, tends to support this assertion (Suppl. Table 2). Increased *Hif1α* expression in particular, is known to significantly increase glycolytic flux while reducing mitochondrial respiration simultaneously limiting mito-ROS and increasing antioxidant activity (59–61). Still, the degree to which a glycolytically biased metabolism and ROS-resilient phenotype pair together along the wound repair timeline and whether these phenotypes are necessary for regeneration in *Acomys* and *Oryctolagus* awaits further testing.

Although we do yet have an explanation for the hyperpolarized mitochondrial phenotype observed in rabbit fibroblasts, future studies using transcriptomic and proteomic screening of the various complex subunits of the electron transport chain may be able to help provide insight. Tracing of Ca^2+^ flux would also be informative in explaining how the membrane potential gradient is seemingly uncoupled from O_2_ consuming capacity, since mitochondrial calcium flux can result in either mitochondrial hyperpolarization (62) or depolarization (63). Assessing mitochondrial pH levels in these cells using dyes like SNARF may provide better resolution on what portion of this observed hyperpolarized state is due to actual ΔpH, and not just increased cation influx into the matrix.

Our study underscores the potential for expanding the use of non-traditional mammalian models to study basal metabolic state and for understanding how tissue repair, inflammation and aging are connected at the molecular level. Moreover, the close phylogenetic proximity of these species has created a paradigm to uncover key genomic, molecular and cellular features that can explain the emergence of regeneration in mammals. Here, two phylogenetically distant mammals share the trait of regenerative ability, coinciding with a shared rapid response to oxidative stress, and finally a shared glycolytic state, which is in turn acquired via different mechanisms involving mitochondrial function and intrinsic mitochondrial ROS production. Further tests screening fibroblasts from multiple mammalian models across different ecosystems and phylogenies would help in creating a mosaic of cellular traits which could support regenerative ability, consisting of different species-specific molecular mechanisms, with proliferative, ROS-adaptive, highly glycolytic, metabolically plastic, low collagen I-producing, non-fibrotic tissue fibroblasts as the pivotal common denominator.

## EXPERIMENTAL PROCEDURES

### Animals

Primary ear pinna cells (see below) were obtained from sexually mature, outbred laboratory mice (*Mus musculus* - ND4 Swiss Webster outbred mice, Inotiv, Strain: Hsd:ND4, RRID:IMSR_ENVIGO:Hsd:ND4, 4-month-old;) and spiny mice (*Acomys dimidiatus,* 4-month-old) bred at the University of Kentucky. While *Mus* were fed 14% protein mouse chow (Tekland Global 2014, Harlan Laboratories, Indianapolis, IN), *Acomys* were fed the same mouse chow supplemented with black-oil sunflower seeds (Pennington Seed Inc., Madison, GA) in a 3:1 ratio. *Mus* and *Acomys* were also given access to water *ad libitum* and maintained on a 14-h light/10-hour dark cycle. Ear tissue from sexually mature (16-week to 20-week-old) New Zealand rabbits (*Oryctolagus cuniculus)* and rats (*Rattus norvegicus* - Sprague-Dawley, Hsd:SD, RRID:RGD_737903) were obtained from S&L Farms (Cynthiana, Kentucky) and from the Division of Laboratory Animal Research, University of Kentucky, respectively. For our aging experiments, primary fibroblasts were obtained from embryonic, young (∼4 mo) and old (>4 yrs) spiny mice. All animals used in our study were female. The University of Kentucky Institutional Animal Care and Use Committee (IACUC) approved all animal work described in this project under protocol 2019-3254.

### Tissue harvest, fibroblast isolation and *in vitro* culture

Primary fibroblasts were obtained from ear pinnae of all animals. Ear pinna tissue was harvested using 4 mm biopsy punches (Sklar Instruments, West Chester, PA) in the centers of the right and left ear pinnae. *Acomys* and *Mus* were anaesthetized with 3% vaporized isoflurane (v/v) (Henry Schein Animal Health, Dublin, OH) at 1 L/min oxygen flow rate while *Rattus* and *Oryctolagus* were euthanized immediately prior to harvest. Tissues from each animal were pooled into separate 1.5 ml Eppendorf tubes. Single-cell suspensions were obtained via enzymatic (Trypsin-Dispase/Collagenase IV) and mechanical digestion (razor dicing/70 µm filtering) steps. The suspension was pelleted and the cell pellets re-suspended and cultured in complete DMEM (Corning® 500 mL Dulbecco’s Modified Eagle’s Medium/Hams F-12 50/50 Mix) supplemented with 10% FBS (Hyclone), and 1% Antibiotic-Antimycotic (Gibco), at 37°C under 3% O_2_ and 5% CO_2_ conditions in 10 cm plates. Cells were cultured and passaged at ∼80% confluency, with each subsequent passage seeded with 300,000 - 500,000 cells in a 10 cm plate. During passages, cells were counted using a hemocytometer.

Fetal fibroblasts were isolated from embryos obtained from timed-pregnant females following euthanasia. The ear anlagen and head skin were carefully dissected from the embryos and subjected to enzymatic digestion in 0.25% trypsin for 10 minutes at 37°C. Following digestion, the enzyme was neutralized using complete growth medium containing serum. Tissue was mechanically dissociated using a 20-gauge needle to obtain a single-cell suspension. Cells were then plated and cultured in Dulbecco’s Modified Eagle Medium/Ham’s F-12 (50/50 mix) supplemented with 10% fetal bovine serum (FBS, HyClone) and 1% Antibiotic-Antimycotic (Gibco). Cultures were maintained at 37°C, under hypoxic conditions (3% O₂, 5% CO₂) in 10 cm tissue culture dishes until reaching confluence. Additional detail on fibroblast isolations can be found here: dx.doi.org/10.17504/protocols.io.yxmvmb3kog3p/v1.

### MitoTracker Staining

Fibroblasts from all species were seeded into 24 well plates on cover slips at 5000 cells per well and cultured for 48 hours in complete DMEM (Corning® 500 mL Dulbecco’s Modified Eagle’s Medium/Hams F-12 50/50 Mix) supplemented with 10% FBS (Hyclone), and 1% Antibiotic-Antimycotic (Gibco), at 37°C under 3% O2 and 5% CO2 conditions. Media was removed and the cells washed with HBSS, and then incubated with MitoTracker Red CMXRos (M7512, ThermoFisher scientific) staining media mix (0.3 µl staining stock/ 1 ml FBS-free DMEM) for 30 minutes. After incubation, cells were washed twice with PBS and then fixed for 15 minutes in the dark with warm 4% formaldehyde. The 4% formaldehyde was removed after fixation and the cells washed twice with PBS for 5 minutes. Chilled acetone was added to the cells at 500 µl/well and incubated for 5 minutes, after which it was removed, and the cells washed once with PBS. DAPI (1:2000/ HBSS) was added to each well and incubated for 15 minutes. The DAPI solution was removed, and the cells finally washed with DI H2O. These coverslips were air dried and mounted on glass slides with Prolong Gold antifade reagent (Invitrogen) and imaged.

For live Mitotracker CMXRos and Mitotracker green (M7514, ThermoFisher scientific) staining, fibroblasts were grown as above. After incubation, the media was removed, washed once with warm 1X HBSS. Before washing, Mitotracker green or Mitotracker CMXRos staining media mix was prepared by mixing 2 µl of the dye stock with 12 ml of FBS-free DMEM and warmed. The cells were incubated in the staining media mix for 30 minutes at the previously mentioned conditions. After incubation, the cells were washed once with 1X HBSS, HBSS added again, and the cells live imaged immediately (12 wells within 20 minutes). Additional detail for this protocol can be found here: dx.doi.org/10.17504/protocols.io.j8nlkyoo6g5r/v1

### Microscopy & Image Acquisition

Manual characterization of mitochondrial phenotypes was carried out using 40X fluorescent images taken with an IX83 microscope (Olympus, Tokyo, Japan) and a DP80 color camera. Images were processed with cellSens software (v1.12, Olympus Corporation, RRID: SCR_014551; https://www.olympuslifescience.com/en/software/cellsens/. 40X images were used for characterizing mitochondrial phenotypes with ∼8 images taken per biological replicate with >30 cells counted per sample. Cells were cultured on coverslips in 24 well plates and fixed before imaging. 20X images were taken for senescence detection with ∼4 images taken per sample and >120 cells counted per sample (n=3/species). Cells positive for individual markers were quantified using ∼5 images captured at 20X magnification per sample, using an Olympus BX53 microscope with each image containing approximately 50 cells. Marker-positive cells were quantified as a percentage of all distinct DAPI positive cells. Live imaging of MitoTracker CMXRos stained cells was carried out using a Nikon CSU-W1 SoRa Confocal Microscope, generating 3D renderings from compilations of z-stacks. Overview images were taken at 60X, and high-resolution images of selected cells were taken at 4 x 60X.

### Mitochondrial morphology analysis

3D stack (nd2 image) files obtained from SoRa microscopy imaging (∼14 per species) were analyzed using the Mitochondrial Analyzer plugin for ImageJ (RRID:SCR_003070; https://imagej.net/). Uploaded images were converted to the 8-bit format, after which respective contrasts were adjusted to reduce background noise. The adjusted images were modified using the 3D Mitochondrial threshold option and the pre-processing commands, “Subtracted background”, “Sigma filter plus” and “Enhance local contrast” were all activated. The post-processing commands “Despeckle”, “Remove outliers” and “Fill 3D holes” were also used. The weighted mean option was selected as the local threshold method, with default block size of 1.250 microns and a C-value of 10. Images taken of *Mus*, *Rattus* and *Oryctolagus* mitochondria were typically thresholded with a block size range of 1-1.250 microns and C-values ranging from 5-10 to prevent the artificial splitting of contiguous mitochondria, since in these cells the mitochondria were many and tubular networks extensive. In *Acomys,* however, because mitochondria were significantly larger and more spheroid block size was set to 4 µm with the C-value at 10. These parameters were defined after using the ImageJ 3D and Nikon NIS Elements viewers to validate that the thresholded images represented the actual 3D projections of the original images. After obtaining consistently representative thresholded images, 3D analysis was carried out on a per cell basis.

### Transmission electron microscopy

Cells were fixed by replacing the growth medium with a fixative solution containing 4% paraformaldehyde and 3.5% EM-grade glutaraldehyde in 0.1 M Sorenson’s phosphate buffer at 4°C. After an initial fixation for 5 minutes at room temperature, the fixative was replaced with fresh solution and incubated at 4°C for 1 hour to ensure complete cross-linking. Following fixation, cells were thoroughly washed with 0.1 M Sorenson’s buffer and then post-fixed in 1% osmium tetroxide for 1 hour at 4°C. Samples were dehydrated in a graded ethanol series 50% through 100% for 5 minutes each, followed by two washes in absolute ethanol, all performed at 4°C. After dehydration, the cells were infiltrated with epoxy resin directly in the culture dish and transferred into beam capsules for overnight embedding. Polymerized blocks were sectioned at 80–90 nm thickness using a Reichert Ultracut E ultramicrotome. Sections were mounted on 300-mesh copper grids and stained with uranyl acetate and lead citrate to enhance contrast. Prepared grids were imaged at the University of Kentucky Electron Microscopy Center using a Talos F200X transmission electron microscope. Additional information for this protocol can be found here: dx.doi.org/10.17504/protocols.io.bp2l6zxxzgqe/v1.

### Fibroblast Oxygen Consumption and Extracellular Acidification Measurement

Measurements of cellular mitochondrial function, and glycolytic flux in fibroblasts were taken with the XF Cell Mito Stress Test (MST) Assay Kit (Agilent Technologies) and XF Glycolysis Stress Test (GST) Kit (Agilent Technologies) respectively, according to the manufacturer’s protocols (https://www.agilent.com/cs/library/usermanuals/public/XF_Cell_Mito_Stress_Test_Kit_User_Guide.pdf; https://www.agilent.com/cs/library/usermanuals/public/XF_Glycolysis_Stress_Test_Kit_User_Guide.pdf) and using the Seahorse Bioscience XFe96 Flux Analyzer (Agilent Technologies). With three biological replicates and three technical replicates for each cell line per species, cells were seeded at passage two into the 96-well Seahorse culture plate at 30,000 – 35,000 cells per well. Glycolytic stress tests, utilizing glucose, Oligomycin and 2-DG, and Mitochondrial stress tests utilizing ETC inhibitors (Oligomycin, FTTP, Rotenone and Antimycin A) were run on live cells 18-24 hours after seeding, per the protocol provided by the manufacturer. From the MST, OCR parameters such as proton leak, maximal respiration, basal respiration, and ATP-linked respiration were quantified, while the GST had ECAR parameters like glycolysis, glycolytic capacity, non-glycolytic acidification, and glycolytic reserve calculated.

### Mass Spectrometry

P2 fibroblasts from *Acomys* and *Mus* (10 million cells per sample) were extracted in 50% acetonitrile (ACN) containing 0.1% formic acid (FA). Following centrifugation, supernatants were collected and analyzed using either an Agilent Infinity II 1290 UHPLC system coupled to a 6546 quadrupole time-of-flight (qTOF) mass spectrometer or an Agilent 1290 Infinity III Bio-UHPLC system coupled to a 6495 triple quadrupole mass spectrometer.

### Untargeted Metabolomics (qTOF-MS)

For untargeted analysis, samples were chromatographed on a 3 µm, 4.6 x 150 mm Luna Omega Polar C18 column. The mobile phase consisted of water containing 0.1% FA (solvent A) and ACN containing 0.1% FA (solvent B). The gradient was as follows at a flow rate of 0.75 mL/min: 0-2 min (15% B), 2-6 min (35% B), 6-8 min (65% B), 8-10 min (100% B), 10-12 min (75% B), 12-14 min (30% B), and 14-18 min (0% B) for re-equilibration. The qTOF-MS was operated in both positive and negative electrospray ionization modes with the following parameters: gas temperature, 300°C; drying gas flow, 12 L/min; nebulizer pressure, 35 psig; sheath gas temperature, 275°C; sheath gas flow, 12 L/min; fragmentor voltage, 100 V; skimmer voltage, 65 V; nozzle voltage, 750 V (0 V in positive mode); and capillary voltage, 3500 V. Data were acquired and processed using Agilent MassHunter Qualitative Analysis, Profinder, and Mass Profiler Professional software.

### Targeted Metabolomics (Triple Quadrupole, MRM Mode)

For targeted analysis, samples were chromatographed on a 2.1 µm, 2.1 x 150 mm Poroshell 120 HILIC-Z column. The mobile phase consisted of ACN (solvent A) and water containing 500 µM ammonium acetate (pH 9.3) and 5 µM medronic acid (solvent B). The gradient was run at a flow rate of 0.3 mL/min as follows: 0-5 min (92% A), 5-10 min (60% A), 10-12 min (40% A), followed by re-equilibration to initial conditions (92% A) from 12-18 min. Quantification was performed using multiple reaction monitoring (MRM) in dynamic MRM mode. For each analyte, two MRM transitions were monitored: a primary transition (quantifier ion) used for quantitation and a secondary transition (qualifier ion) used for confirmation of analyte identity. Analyte identification was based on retention time matching with authentic standards and consistency of the quantifier-to-qualifier ion ratio. Collision energies voltages were optimized individually for each analyte using standard solutions. Deuterated salicylic acid and benzocaine were used as internal standards to correct for extraction efficiency, ionization variability, and instrument drift. Data were acquired and processed using Agilent MassHunter - Qualitative and Quantitative Analysis software.

### RNA-seq read processing, expression quantification, and differential expression analysis

Bulk RNA was performed on primary fibroblast samples from *Acomys dimidiatus* and *Mus musculus* to generate paired-end, 150 bp poly(A)-enriched libraries on the DNB Eukaryotic Strand-Specific RNA-seq platform (mean counts / sample ∼29 million). Read quality was assessed with FastQC and summarized with MultiQC (64). Upstream raw read processing followed a conventional alignment-based bulk RNA-seq workflow (65). Reads were aligned to species-specific reference genomes with HISAT2 (66), converted and sorted to BAM format with SAMtools (67), and quantified against reference annotation gene models with StringTie (68). Transcript assembly and abundance estimation were performed using StringTie in reference-guided mode. Gene-level count matrices were generated using the prepDE.py3 script from the StringTie workflow for importing into downstream analysis. Cross-species gene comparisons were restricted to a subset of conserved, strictly one-to-one orthologs. Ortholog relationships were identified with Orthofinder2 (69) (v2.5.4) using Multiple Sequence Alignment (MSA) mode with the *Acomys dimidiatus* GCA_907164435.1 Ensembl rapid release and the *Mus musculus* GRCm38 (v102) Ensembl genome annotations. These results were trimmed into a shared one-to-one subset anchored to *Mus* Ensembl gene identifiers that exclude paralogs or multiple alignments. Gene IDs were keyed to a stable *Mus musculus* Ensembl gene identifier and gene symbol using a local Ensembl GRCm39 export. The trimmed transcriptome contained 12,944 ortholog-aligned genes after intersection with the full six-sample count matrix. Differential expression was modeled with DESeq2 (70) using a Wald test with a model ‘∼ species’ on the one-to-one ortholog count matrix, with *Acomys dimidiatus* as the reference level. *P*-values were adjusted with the standard Benjamini-Hochberg procedure, and genes with FDR < 0.05 were considered significantly differentially expressed. Shrunken log2 fold changes were estimated with apeglm for effect-size reporting (71), while the raw DESeq2 adjusted p-values were used for significance calls.

### Curated Heatmap

To visualize gene expression levels between species, heatmaps were generated using ComplexHeatmap (72) with curated panels of significantly, differentially expressed genes for glycolysis/gluconeogenesis, pyruvate metabolism, Krebs/TCA metabolism, glutathione synthesis, import and cycling. For each gene, the DESeq2-normalized count was transformed as log2(normalized count + 1), then z-normalized across the six samples. These row z-scores are used to display within-gene expression patterns across samples in the heatmap. Genes are hierarchically clustered within each curated block, while sample and block order are manually fixed for visualization.

### Mitochondrial membrane potential characterization: JC-1 Staining

P2 fibroblasts from *Acomys*, *Rattus*, *Mus* and *Oryctolagus* were plated on a 96-well black clear glass bottom plate and incubated for 24-48 hours. This dye is also cationic and labels mitochondria with respect to mitochondrial membrane potential reversibly. The JC-1 dye, which is monomeric, specifically binds to all mitochondria and has a green fluorescence emission of ∼529nm. Upon encountering “healthy” mitochondria with high membrane potential, the cationic dye accumulates in the mitochondrial matrix and forms aggregates, with a red fluorescence emission at ∼590nm. The red/green fluorescence intensity ratios are then used to describe mitochondrial polarization state. The JC-1 dye is typically used as an indicator of mitochondrial health in response to different chemical stimuli, hinged on the principle that mitochondrial membrane potential disruption results in dysfunctional mitochondria, signifying early tell-tale signs of apoptosis. Here, this was used to compare mitochondrial polarization levels across the four species. For basal assessment of mitochondrial polarization, fibroblasts from the four species (4 biological replicates per species) were cultured on glass bottomed dark 24-well plates till they reached ∼80% confluency, incubated with the dye for 30 minutes, washed and live-imaged. Fluorescence intensity in the red and green channels was quantified in the images taken using ImageJ.

To measure changes in mitochondrial membrane potential in response to mitochondrial inhibitors FCCP, and Oligomycin, cells were seeded on a 96-well black clear bottom plate. Cells were plated at a density of ∼30,000 cells per well and incubated for 24 hours. During plating, allowance was made for controls and two treatment conditions. Cells were washed in 1X HBSS after which a staining media mix (2µM) was prepared and added at 100µl per well for 30 minutes. Halfway through the incubation period 1.5µM of Oligomycin and 10mM of FCCP were added to the treatment wells to complete the incubation. After a wash with warm 1X HBSS, the plates were scanned with excitation set at 485nm and emission detection set at 529nm for the green monomeric form and 590nm for the red aggregate form. Three washes with three corresponding readings were carried out. The ratio of green to red fluorescence signal intensities were used to quantify the levels of mitochondrial membrane potential. The values were normalized to ug of sample protein. Additional protocol detail can be found here: dx.doi.org/10.17504/protocols.io.rm7vz9xxrgx1/v1.

### Mitochondrial Isolation

Cultured fibroblasts were dislodged with trypsin (at P2), spun down (1500 rpm for 5 minutes) five minutes after the reaction was stopped with FBS-containing DMEM. The supernatant was poured off and pellets kept on ice. 1 ml of Mitochondria Isolation Buffer (MIB) was added to the pellet and the pellet re-suspended. The suspension was spun at 400 rcf for 5 minutes, after which the supernatant was taken out and 450 µl of EDTA added to the pellet and re-suspended. Samples were broken up using a Nitrogen cell disruptor (1250psi for 10 minutes). The samples cell lysates were passed twice through a 26-gauge syringe, topped up with 1.5 ml of Mitochondria Isolation Buffer, spun at 1300 x *g* for 3 minutes at 4°C and the supernatant transferred into new tubes. These were then spun at 4°C at 1300 x *g* for 10 minutes. A Ficoll gradient was made up using 25 ml of STE (8.55g of sucrose, 32 µl of 0.5 M EDTA, 80 µl of 1 M Tris and topped up to 25 ml with dIH_2_O) and a Ficoll stock (10 ml STE and 17.5 ml 20% Ficoll). The mitochondria were added to and then spun down in the Ficoll gradient (2 ml 7.5%/ 2 ml 10%). After spinning at 32,000 rpm for 30 minutes, the Ficoll supernatant was discarded, and the pellet resuspended in 1 ml MIB and spun at 13,000 rcf for 10 minutes. The supernatant was then discarded and the pellets (purified mitochondria) resuspended in MIB and quantified using a BSA assay. Additional protocol detail can be found here: dx.doi.org/10.17504/protocols.io.81wgbwxr1gpk/v1.

### Electron Transport Chain Protocol (ETCP)

A modified version of the Respirometry in frozen samples (RIFS) assay (40) called the Electron transport chain protocol (ETCP) (39) was carried out using n=4 biological replicates/species (aka four independently derived primary lines per species): i.e., frozen mitochondria previously isolated from cultured *Acomys*, *Mus*, *Rattus* and *Oryctolagus* ear pinna fibroblasts (P2). After thawing the samples, 25µl of Isolation buffer was added to all samples and gently resuspended. 4µl of mitochondrial mix was loaded per well (2 technical replicates) and the protein concentration for each sample was measured using the BCA assay. Seahorse XF 96 well cartridge ports were prepared as follows: Port A (ETC 1): 3220µl Mitochondrial Respiration buffer w/o BSA, 2.8µl Rotenone, 280µl succinate; Port B: 3500µl Mitochondrial Respiration buffer w/o BSA, 3.5µl Antimycin A; Port C: 3325µl Mitochondrial Respiration buffer w/o BSA, 88µl Ascorbate, 88µl TMPD; Port D: 3033µl Mitochondrial Respiration buffer w/o BSA, 480µl Azide. Each well had each port loaded with 25µl of the port mixes. With the Seahorse XF 96 cell culture plate on ice, 75µl mitochondrial samples (2µg protein) were loaded per well, after which the plate was balanced and spun at 3214 rcf for 10 minutes at 4°C. 11ml of the Alamethicin Assay solution (10.6ml Mitochondrial Respiration buffer w/o BSA, 40mg/ml Alamethicin 9.63µl, 1.5mM NADH 385µl, 10 µM cytochrome c 48µl) was prepared with 100µl of the assay solution added to each well and ran on the Seahorse XF 96 Analyzer immediately. With the contents of Ports A-D injected sequentially over a period of 60 minutes, Complex 1, 2 and 4 activities were determined. Additional protocol detail can be found here: dx.doi.org/10.17504/protocols.io.6qpvrw39olmk/v1.

### Statistical analyses

All *in vitro* experiments except the aged fibroblast panel used n=4 biologically independent samples (cell lines from separate animals) and a minimum of 3 technical replications. The lifespan panel and bulk RNA-seq analysis used n=3 biologically independent lines. We did not calculate power. Different animals were used across assays and experiments such that a total of ∼8-12 animals/species were used for this study. Data were initially compiled using Microsoft Excel (RRID:SCR_016137; https://www.microsoft.com/) with JMP (version Pro 17.1.0, SAS Institute Inc., https://www.jmp.com/en/software/data-analysis-software, RRID:SCR_014242) used to perform all statistical analyses. Data from Seahorse assay, respirometry, fluorescence signal intensity experiments were presented as mean ± SD and had their statistical significance determined using one-tailed or two-tailed ANOVA with all post hoc pairwise comparisons made with the Student’s *t*-test. Alpha was set at 0.05 to assess significance for all statistical analyses.

## Supporting information

Revised Supp. Data and Tables

## ACKNOWLEDGEMENTS

We would like to thank Josh Sarli, Brennan Riddell and Ava Musarra for animal husbandry. We thank all members of the Seifert lab for helpful discussions in developing the manuscript. This research was funded in part by grants from the NIH (R01AR070313) and Aligning Science Across Parkinson’s (ASAP-020495) through the Michael J. Fox Foundation for Parkinson’s Research (MJFF) to AWS, VA Merit Award (2I01BX003405) (PGS), NIH (R01 NS112693-01A1) (PGS) and NIH (P20 GM148326-01) (PGS) and USDA National Institute of Food and Agriculture (Grants-13323564 and 14138983 and Hatch project 1014539) (PK). For the purpose of open access, the author has applied a CC BY public copyright license to all Author Accepted Manuscripts arising from this submission. Access to characterization instruments and staff assistance was provided by the Electron Microscopy Center at the University of Kentucky, member of the KY INBRE (Kentucky IDeA Networks of Biomedical Research Excellence), which is funded by the National Institutes of Health (NIH) National Institute of General Medical Science (IDeA Grant P20GM103436). Chemical analysis reported in this study was carried out at the Center for Agricultural and Life Sciences Metabolomics (CALM; https://plantpathology.ca.uky.edu/lab/Analytical-CORE).

## AUTHOR CONTRIBUTIONS

**Ebenezer Aryee:** Conceptualization, Methodology, Visualization, Formal Analysis, Investigation, Data Curation, Writing-original draft preparation, reviewing and editing. **Sandeep Saxena**: Methodology, Investigation. **Renee Donahue**: Methodology. **Brennan Riddell:** Methodology, Formal Analysis, Investigation, Data Curation. **Hemendra Vekaria**: Methodology. **Patrick Sullivan**: Methodology, Resources. **Kachroo Pradeep:** Methodology, Resources. **Huazhen Liu:** Methodology. **Ajoy Aloysius**: Methodology, Investigation. **Ashley Seifert:** Conceptualization, Methodology, Investigation, Writing-original draft preparation, review and editing, Funding acquisition, Project administration, Resources, Supervision. **All authors**: Writing - review and editing.

## DATA AVAILABILITY

The code, as well as data, protocols, and key lab materials used and generated in this study are listed in a Key Resource Table alongside their persistent identifiers (Supplementary Data and Tables) which is available on the Zenodo platform via the ASAP Data repository with DOIs: https://doi.org/10.5281/zenodo.23269045 and https://doi.org/10.5281/zenodo.23270260 respectively. In addition, the entire mitochondrial imaging dataset consisting of all images from live MTR, MTG, JC-1 stains as well as images from fixed cells (TEM and MTR-stained), have been made available on the Zenodo platform via the ASAP Data repository: https://doi.org/10.5281/zenodo.21416619.

## Supplementary Figures

**Supplementary Figure S1.**
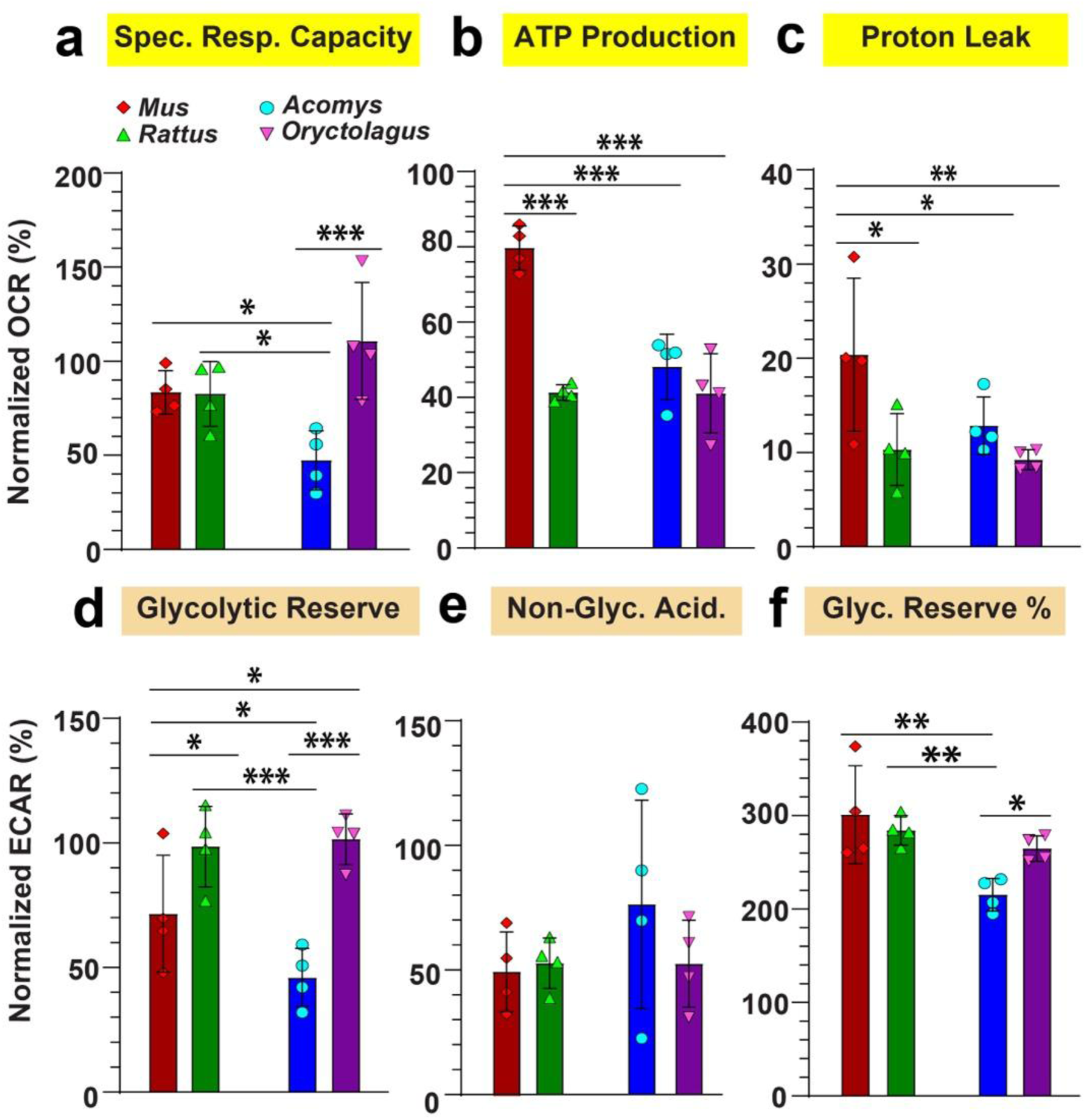
Glycolytic and mitochondrial respiration parameters in *Mus*, *Acomys Rattus* and *Oryctolagus* fibroblasts. Oxidative Phosphorylation and glycolytic parameters determined in all four species from the MST-mitochondrial stress tests **(a-c)** and GST-Glycolytic stress tests **(d-f)** run on Seahorse XFe96 analyzer using primary ear pinna fibroblasts (continued from Fig. 1). While *Mus* showed the highest levels of ATP production (ANOVA, *F*=24.0693, *p*<0.0001) and proton leak (ANOVA, *F*=4.4067, *p*=0.0262), rabbit fibroblasts interestingly shared higher specific respiratory capacity with *Mus* and *Rattus*, all three significantly higher than *Acomys* (ANOVA, *F*=6.5691, *p*=0.0071). Glycolytic reserve levels were distinctively higher in *Rattus* and *Oryctolagus* compared to *Acomys* and *Mus* (ANOVA, *F*=10.3251, *p*=0.0012). All ECAR parameters across all four species were normalized to the average basal glycolytic rate in *Mus*. All OCR parameters were also normalized to the average basal respiration rate in *Mus*. Individual data points represent biological replicates (n=4) for each species. Post-hoc pairwise comparisons were carried out using the Student’s *t*-test. (* *p*<0.05, ** *p*<0.001, *** *p*<0.0001).

**Supplementary Figure S2.**
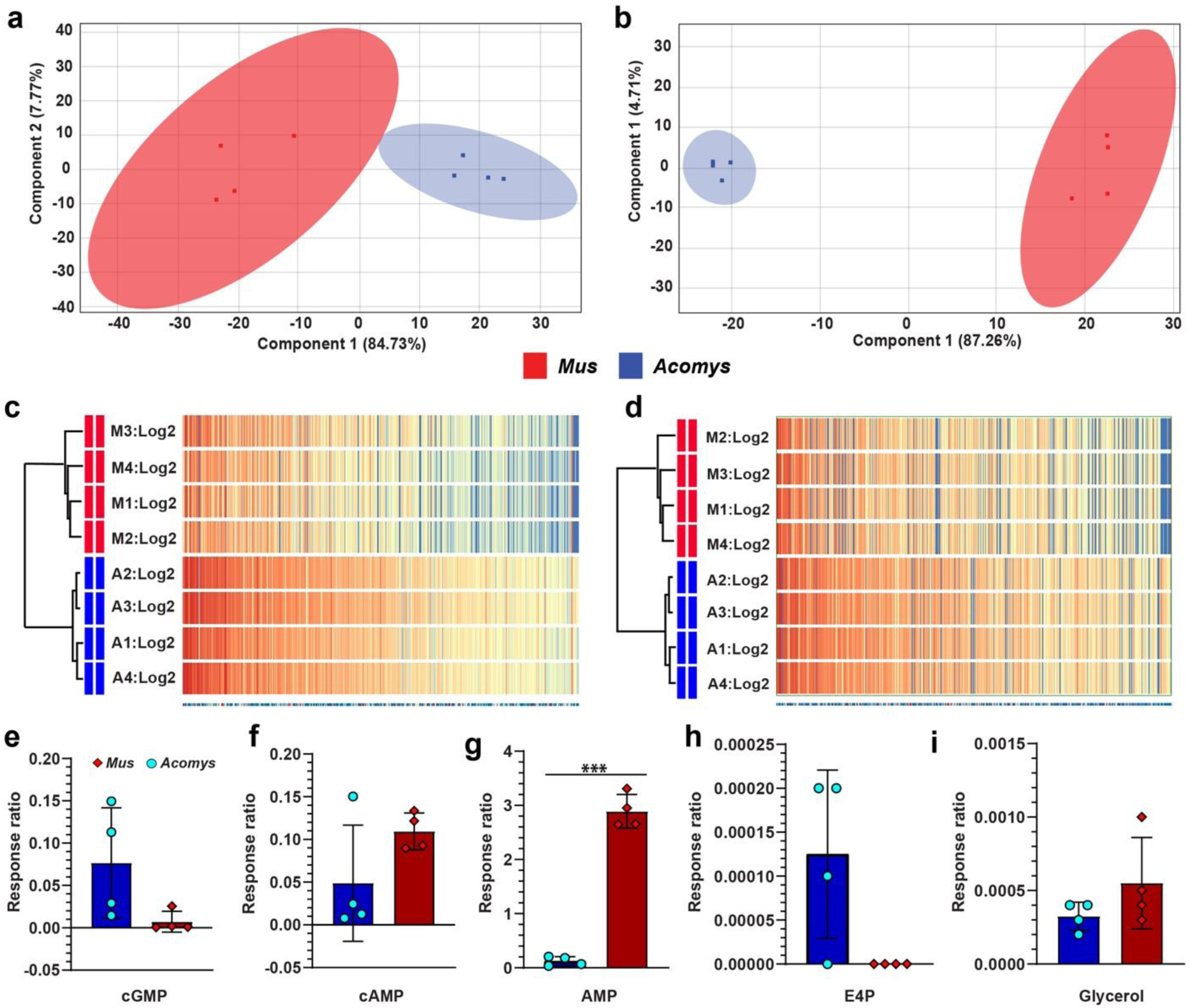
Metabolomic comparison of *Acomys* and *Mus* fibroblasts. (**a-d**) Untargeted metabolite profiling indicates that *Mus* and *Acomys* fibroblasts (4 biological replicates/species) are cluster distinctly when positively charged (b,d) and negatively charged (a,c) compound analysis is carried out. (**e-i**) Targeted metabolomic analysis of energy-associated metabolites in *Acomys* and *Mus* fibroblasts. Pairwise comparisons were carried out using the Student’s *t*-test. (*** *p*<0.0001).

**Supplementary Figure S3.**
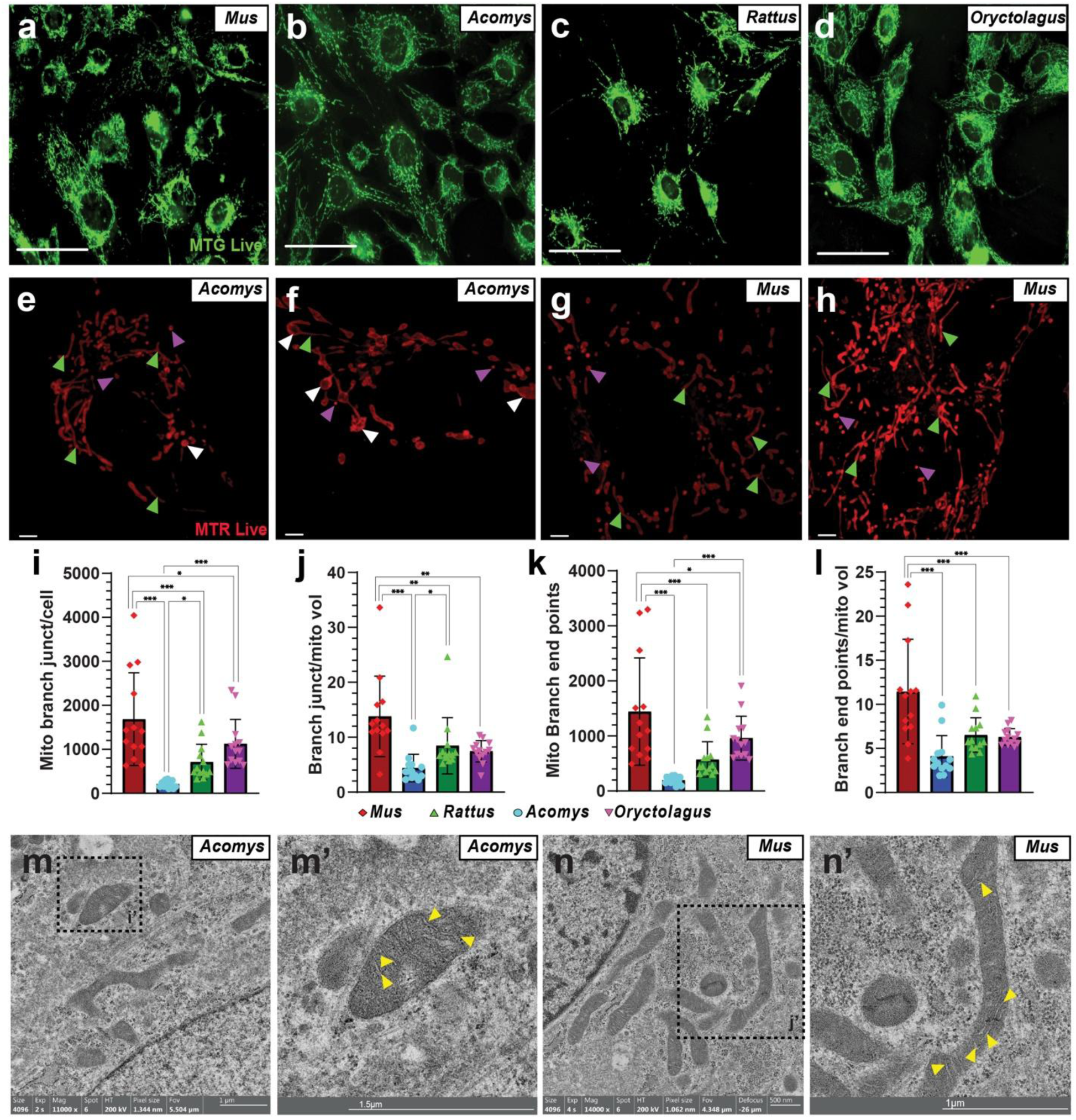
Mitochondrial characterization in primary fibroblasts. (**a-d**) Mitochondrial characterization using MitoTracker green staining and live imaging revealed tubular mitochondrial networks in all four species (Scale bars: 20µm). Mitochondrial density in *Acomys* was comparatively lower with distinctive round structures, although not as massive as seen in MTR-stained cells (b). (**e-h**) Primary *Mus* and *Acomys* fibroblasts stained with MTR and live imaged via spinning disc confocal microscopy (Scale bars: 5µm). *Acomys* showed a range of features, one with more tubular mitochondria (e) (green arrows), others with more massive spherical mitochondria (f) (white arrows), although the massive spherical phenotype was dominant. Purple arrows indicate small, circular mitochondria. Tubular mitochondrial networks were generally consistent in *Mus* (g-h). (**i-l**) Mitochondrial analysis of 3D images indicate differences in the number of mitochondrial branch junctions per cell (ANOVA, *F*= 12.7996, *p*<0.0001), number of branch junction per mitochondrial volume (ANOVA, *F*= 8.8596, *p*<0.0001), number of mitochondrial branch end points (ANOVA, *F*= 12.1688, *p*<0.0001), and number of mitochondrial branch end points per total mitochondrial volume (ANOVA, *F*= 11.0055, *p*<0.0001) in the four species. (**m-n)** TEM images of fibroblasts obtained from young *Mus* and *Acomys*. Yellow arrows indicate cristae.

**Supplementary Figure S4.**
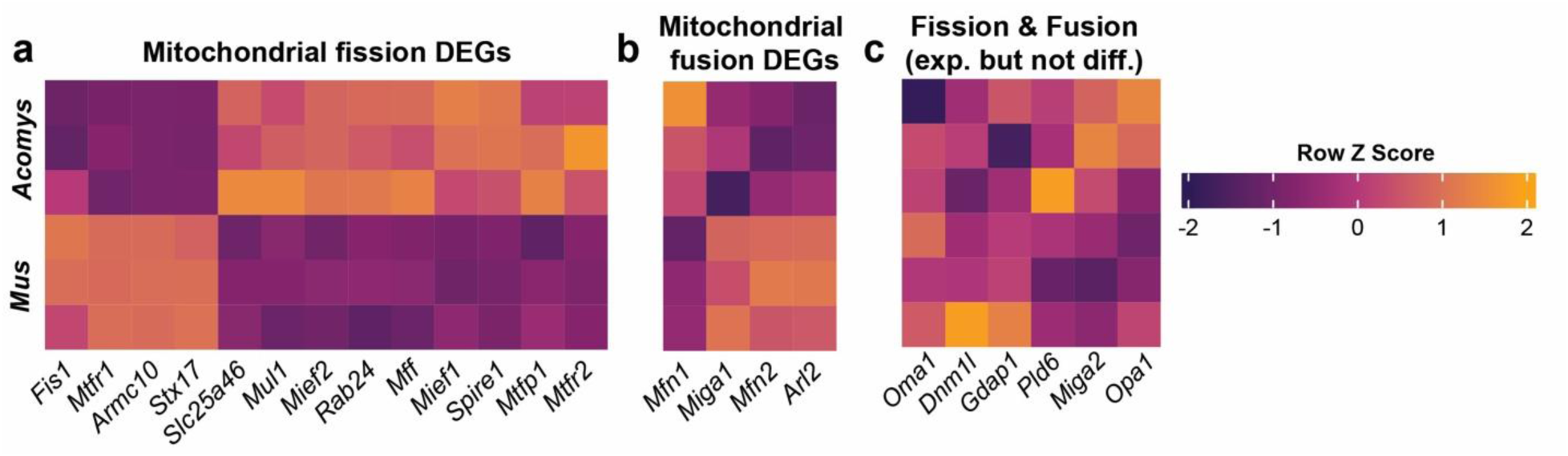
Mitochondrial fusion and fission gene expression in *Acomys* and *Mus* fibroblasts does not correlate with observed differences in mitochondrial morphology. (**a-c**) Heatmaps generated from bulk RNA-seq comparison of *Acomys* and *Mus* fibroblasts (n=3/species) showing DEGs of key enzymes involved in mitochondrial fission (a), DEGs of key enzymes involved in mitochondrial fusion (b), and critical genes involved in mitochondrial fission and fusion which are expressed in both species but whose expression patterns do not significantly differ (c). All genes included in heatmaps (a-b) are significantly-differentially expressed across biological replicates (n=3) between the two species (FDR<0.05).

